# Oligodendroglial dysfunction drives parvalbumin interneuron hypomyelination, circuit dysfunction and social deficits in an ASD model

**DOI:** 10.64898/2026.11.12.766147

**Authors:** M Villafranca-Faus, I Ugarte-Arakistain, A Sanchez-Camacho, C Peiró-Moreno, A Robledo, S Calovi, P Mata, L Bayon-Cordero, O Fresnedo, MV Sanchez-Gomez, Y Rueda, A Battefeld, FN Soria, O Peñagarikano, JM Rosa, M Domercq

## Abstract

Altered social behaviour in autism spectrum disorders (ASDs) has been linked to disruptions in neural circuit development, with emerging evidence implicating abnormal myelination. Yet how myelin development is affected and how this impacts circuit function remain unclear. Here, using the *cntnap2^-/-^* mouse model of ASD, we show that oligodendrocyte precursor cell (OPC) differentiation is transiently impaired during early postnatal development, resulting in delayed myelination in the somatosensory cortex and corpus callosum, altered myelin lipid composition, and impaired sheath wrapping, accompanied by transient microgliosis and increased oligodendrocyte apoptosis. Transcriptomic analysis reveals upregulation of interferon and immune-related pathways alongside downregulation of GABAergic synaptic genes in oligodendrocytes. These alterations converge on disrupted GABAergic interneuron–OPC communication, leading to hypomyelination and impaired maturation of parvalbumin-positive (PV⁺) interneurons through a non-cell-autonomous mechanism, with selective deficits in PV⁺ function, excitation–inhibition balance and gamma oscillations. Early treatment with the promyelinating agents clemastine or pranlukast restores OPC differentiation, myelination, and PV⁺ maturation, rescues circuit dysfunction, and ameliorates social deficits and behavioural inflexibility. Our findings establish that transient developmental failure in OPC differentiation drives PV⁺ interneuron hypomyelination, linking oligodendroglial dysfunction to circuit-level and behavioural alterations in ASD, and highlight early myelin-targeted interventions as a potential therapeutic strategy.

## INTRODUCTION

Autism spectrum disorder (ASD) is a clinically heterogeneous neurodevelopmental condition characterized by deficits in social communication and interaction, restricted and repetitive behaviours, and atypical sensory processing^1^. Typically diagnosed in early childhood, ASD affects an estimated 1% of the population worldwide and up to 2.8% in the United States^2^, and is associated with enduring cognitive, behavioural, and physiological impairments across the lifespan^1^. ASD has a strong but heterogeneous genetic basis, with contributions from both genetic and non-genetic factors. Genome-wide association and sequencing studies have identified hundreds of risk genes implicated in diverse biological processes, including transcriptional regulation, synapse development, neurotransmission, and activity-dependent plasticity^3–5^.

Despite this genetic complexity, converging evidence points to shared circuit-level mechanisms underlying ASD pathophysiology. Among the most influential frameworks is the excitation/inhibition (E/I) imbalance hypothesis, which posits that disrupted inhibitory control, largely due to GABAergic interneuron dysfunction, leads to aberrant network dynamics and impaired information processing^6,7^. Within inhibitory circuits, fast-spiking parvalbumin-expressing (PV⁺) interneurons have emerged as a particularly critical cellular substrate. PV⁺ interneurons provide powerful perisomatic inhibition onto pyramidal neurons and are essential for the generation of gamma oscillations, sharp-wave ripples, and precise spike timing-network dynamics fundamental to learning, memory, and cognitive flexibility^8,9^. Notably, PV⁺ interneuron maturation is a protracted postnatal process driven by activity-dependent transcriptional programs, including PGC-1α^10^, and requiring the establishment of perineuronal nets and integration of glial signals^11,12^. Reduced PV expression has been consistently reported in postmortem cortical tissue from individuals with ASD^13,14^. Mouse models carrying loss-of-function variants in ASD-associated genes, including *Shank3, Mecp2*, and *cntnap2*, similarly exhibit reduced PV levels, impaired inhibitory synaptic transmission, and disrupted network oscillations^13,15–19^, though the mechanisms underlying PV downregulation remain unclear.

Beyond synaptic dysfunction, increasing evidence implicates abnormal brain connectivity and myelination deficits as key contributors to ASD pathophysiology. Integrative transcriptomic analyses across multiple ASD mouse models and human postmortem samples have identified consistent dysregulation of oligodendrocyte lineage genes and myelin-related pathways^20^, suggesting that altered myelination may represent a shared pathological feature of ASD^21–25^. Myelination is essential for regulating axonal conduction velocity, temporal precision, and synchrony of neuronal firing—properties critical for efficient network communication and higher cognitive functions^26^. Over the past decade, activity-dependent or adaptive myelination has emerged as a key mechanism for fine-tuning neural circuit function during development and adulthood^26–28^, supporting motor learning, memory consolidation, and social behaviour^28–31^. Critically, myelination also plays a direct role in PV⁺ interneuron maturation. GABAergic signalling between PV⁺ interneurons and oligodendrocyte precursor cells (OPCs) regulates activity-dependent myelination^32,33^, which in turn stabilizes fast-spiking properties and inhibitory efficacy.

Disruption of this bidirectional communication leads to impaired PV firing, hypersynchronous network activity, and deficits in cognition and social behaviour^32–34^. These findings raise the possibility that myelination deficits may represent an upstream mechanism contributing to PV⁺ interneuron dysfunction and E/I imbalance in ASD.

Despite converging evidence linking E/I imbalance, PV⁺ interneuron dysfunction, and myelination abnormalities, the causal relationships between these processes remain poorly understood. Here, we address this gap by investigating whether altered myelination contributes to PV⁺ interneuron dysfunction during development in an ASD model. Cntnap2, classified as a high-confidence ASD risk gene by the Simons Foundation Autism Research Initiative (SFARI), encodes the axonal adhesion molecule Caspr2^35^. Loss-of-function mutations in *cntnap2* identified in ASD patients^36^ and *cntnap2* polymorphisms^37^ have been associated with increased ASD risk. *Cntnap2*^-/-^ mice recapitulate key ASD-related phenotypes, including epileptiform discharges, hyperactivity, social interaction deficits, and repetitive behaviour^15^. Using this model, we show that OPC differentiation and myelination are transiently impaired during early postnatal development, with a pronounced impact on PV⁺ interneuron maturation, function, and circuit dynamics. Importantly, early promotion of OPC differentiation restores PV⁺ interneuron myelination, maturation and function, normalizes E/I balance and gamma oscillations, and rescues social behaviour and cognitive flexibility. Together, these findings identify developmental myelination as a critical regulator of inhibitory circuit maturation and a potential therapeutic target in ASD.

## RESULTS

### Early delay in myelination and oligodendrocyte differentiation in *cntnap2*^-/-^ mice

Although myelin abnormalities have been reported in several animal models of ASD^20–25^, their developmental dynamics have not been systematically examined. Relative to other cellular developmental processes in the brain, oligodendrogenesis and myelination are remarkably protracted, extending across the first postnatal weeks in mice. We therefore analyzed cortical myelination in the somatosensory cortex (S1) at postnatal day 20 (P20, juvenile) and postnatal day 60 (P60, adult) in wild-type (WT) and *cntnap2*^-/-^ mice. We focused on S1 because it plays a key role in sensory processing, which is frequently altered in ASD and is closely linked to core behavioural features. In juvenile *cntnap2*^-/-^ mice, we observed a significant delay in myelination, as evidenced by a reduction in the proportion of the cerebral cortex myelinated, which was nearly restored in adults (Fig. 1A). Within layer V, myelin coverage (length) was reduced in juvenile *cntnap2^-/-^* mice but restored in adults (Fig. 1B). Accordingly, the number of Nav1.6⁺ nodes, flanked by Caspr⁺ paranodes, was significantly reduced in layer V in juvenile *cntnap2*^-/-^ mice (Fig. 1C) but practically restored in adults. Finally, transmission electron microscopy revealed a significant increase in g-ratio in juvenile *cntnap2*^-/-^ mice (Fig. 1D). Although myelination was nearly restored in adults, a more detailed analysis of the columnar pattern of myelination revealed an altered organization and orientation of myelinated fibres that persisted in adult *cntnap2*^-/-^ mice (Fig. 1E). Myelin deficits were not exclusive to S1. Similar deficits in myelination were detected in the corpus callosum of juvenile *cntnap2*^-/-^ mice. The density of myelinated fibres as well as the number of nodes of Ranvier were significantly reduced in juvenile *cntnap2*^-/-^ mice and nearly restored in adults (Fig. S1A, B). No differences were detected in node length (Fig. S1B). TEM analysis showed a significant increase in g-ratio in juvenile *cntnap2*^-/-^ mice (Fig. S1C). Compound action potential (CAP) recordings from the corpus callosum did not reveal any significant change in nerve conduction velocity of myelinated (N1) or unmyelinated (N2) fibres in juvenile *cntnap2*^-/-^ mice (Fig. S1D). However, the amplitudes of the N1 and N2 peaks were altered and the N1/N2 ratio was reduced in *cntnap2*^-/-^ mice, further supporting a decrease in the proportion of myelinated fibres and an increase in unmyelinated ones (Fig. S1D). In summary, our data point to an impairment of myelination during development and to subtle changes in myelin patterns in adult *cntnap2*^-/-^ mice.

**Figure 1.**
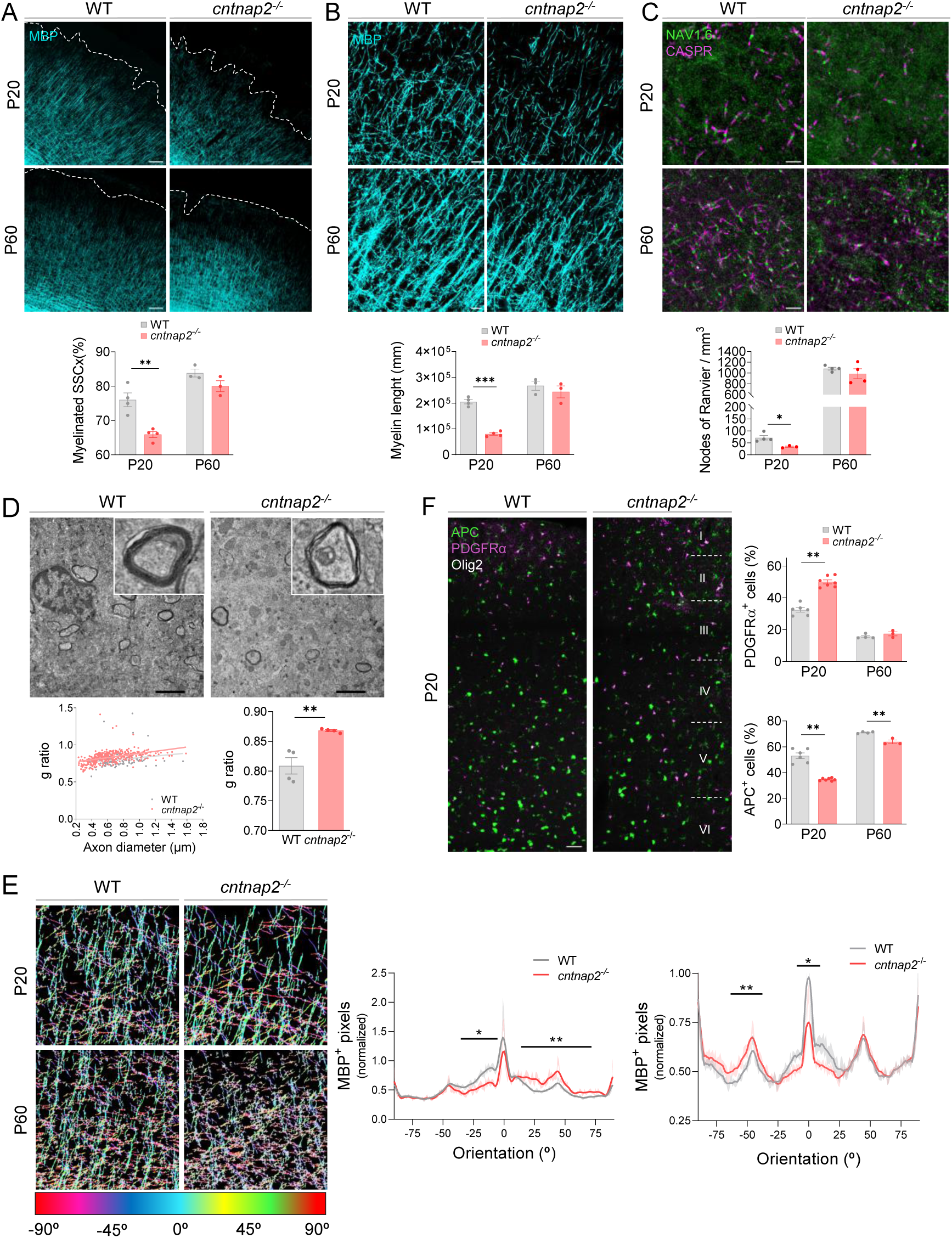
Impairment of myelination during development in the somatosensory cortex (S1) of *cntnap2*^-/-^mice. **A-C**) Confocal z stack images displaying MBP staining (A, B) and sodium channels Nav1.6/ Caspr labeling of nodes of Ranvier (C). Scale bar = 100 μm (A), 20 μm (B) and 5 μm (C). Quantification of myelinated area (A, total S1), myelin coverage (B, layer V) and the number of nodes of Ranvier (C, layer V) in juvenile (P20) and in the adult (P60) WT and *cntnap2*^-/-^ mice (n = 4/4 at P=20 and n=3/3 at P60 for WT and *cntnap2*^-/-^ mice respectively). **D**) TEM images of myelinated axons in layer V of the S1 in juvenile WT and *cntnap2*^-/-^ mice. Scale bar = 2 μm. Below, g-ratio distribution (left) and average (right) in juvenile WT and *cntnap2*^-/-^ mice (n=4/4). **E**) Myelinated fiber orientation analysis showed that columns pattern of myelination is altered in both juvenile and adult *cntnap2*^-/-^ mice (n=4/4 at P20 and 3/3 at P60). **F**) Confocal z stack images displaying triple staining, APC (oligodendrocytes), PDGFRα (OPCs) and olig2 (all the oligodendroglial lineage) in S1 of juvenile WT and *cntnap2*^-/-^ mice. Scale bar = 50 μm. *Rigth*, proportion of PDGFRα^+^ OPCs and APC^+^ mature oligodendrocytes normalized to the total number of olig2^+^ oligodendrocytes in juvenile and adult WT and *cntnap2*^-/-^ mice (n= 4/4 at P20 and 3/3 at P60). Data are represented as mean + SEM. Statistical analysis were performed with 2-tailed unpaired Student t-test. *p<0.05, **p<0.01, ***p<0.001.

To assess whether these defects reflect altered oligodendrocyte differentiation, we performed triple immunostaining for PDGFRα (OPCs), APC (mature oligodendrocytes), and Olig2 (pan-oligodendroglial marker). In S1 of juvenile *cntnap2*^-/-^ mice, the proportion of mature APC⁺ oligodendrocytes was significantly reduced, while the number of PDGFRα⁺ OPCs was increased (Fig. 1F). Although lineage progression tended to normalize in adults, a significant decrease in mature APC⁺ oligodendrocytes in S1 persisted (Fig. 1F). In the corpus callosum, a similar delay in oligodendrocyte differentiation was detected in juvenile *cntnap2*^-/-^ mice, and in this case adult mice recovered completely (Fig. S1E). In both regions, the number of BCAS1⁺ immature oligodendrocytes, representing an intermediate differentiation state, tended to be increased at P20 (Fig. S2). Altogether, these results indicate that myelin deficits during development in *cntnap2*^-/-^ mice tend to normalize in the adult.

### Non-cell-autonomous oligodendrocyte dysfunction is associated with early immune dysregulation in *cntnap2^-/-^* mice

To determine whether the myelination deficits observed in vivo reflect cell-autonomous oligodendrocyte dysfunction, we assessed differentiation and myelination capacity of *cntnap2*^-/-^ oligodendrocytes *in vitro*. The proportions of MBP⁺ mature oligodendrocytes and NG2⁺ OPCs were comparable between genotypes, and myelination of nanofibers was unaffected (Fig. 2A), demonstrating that *cntnap2*^-/-^ oligodendrocytes retain full intrinsic differentiation and myelination capacity. Consistent with this, publicly available transcriptomic databases confirmed negligible *cntnap2* expression in the oligodendrocyte lineage. These findings indicate that the myelination deficits in *cntnap2*^-/-^ mice arise non-cell-autonomously, reflecting the response of oligodendrocytes to an altered CNS microenvironment.

**Figure 2.**
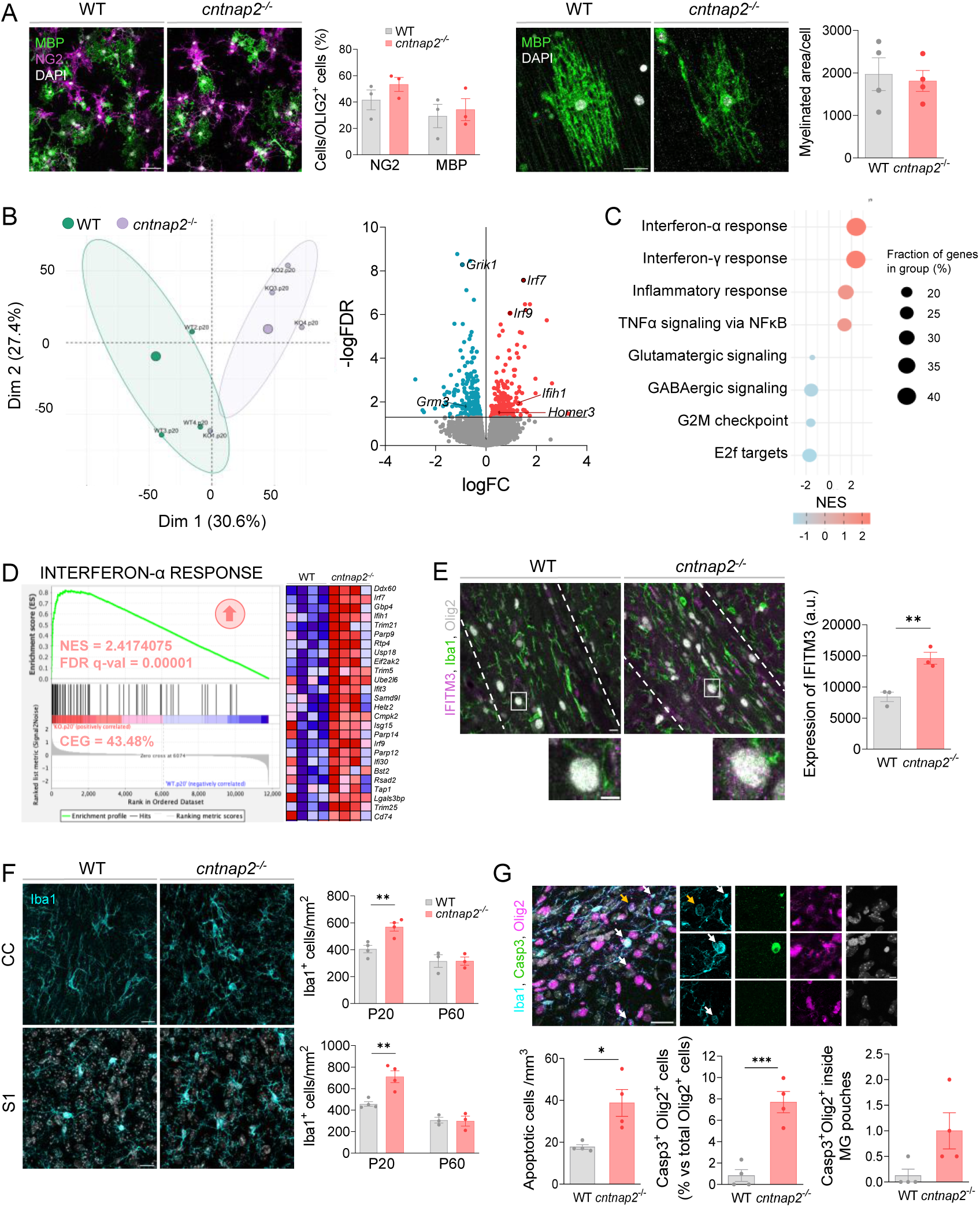
Transcriptional profiling identified immune signaling upregulation in juvenile *cntnap2*^-/-^ oligodendrocytes. **A**) Proportion of NG2^+^ OPCs and APC^+^ oligodendrocytes as well as nanofiber myelination in vitro from WT and *cntnap2*^-/-^ mice (n= 4 different cultures). **B**) Left, principal components analysis (PCA) of WT and *cntnap2*^-/-^ oligodendrocytes at P20. Right, volcano plot showing differentially expressed genes (DEGs, FDR q-value <0.05) in juvenile *cntnap2*^-/-^ mice. Upregulated genes (210) are shown in red and downregulated genes (234) in blue. Relevant DEGs are indicated. **C**) Hallmark pathway analysis of RNA sequencing data with Gene Set Enrichment Analysis (GSEA) showing top 4 upregulated (red) and downregulated (blue) biological processes, ranked by normalized enrichment score (NES). **D**) Enrichment plots (left) of Interferon gene sets from hallmark analysis with corresponding NES, FDR q-value and Core Enrichment Genes (CEG). CEG is defined as the percentage of significant genes within the set. On the right, heatmap of the most significant genes. **E**) Confocal z stack images and quantifications of IFITM3 expression in oligodendrocytes from juvenile WT and *cntnap2^-/-^* mice (n = 3). **F**) Confocal z stack images at P20 and quantifications at P20 and P60 of Iba1^+^ microglia cells in corpus callosum and S1 of WT and *cntnap2*^-/-^ mice (n=4/4 at P20 and 3/3 at P60). Scale bar = 15 μm. **G**) Confocal z stack images showing Cap3^+^ (green) Olig2^+^ (red) apoptotic oligodendrocytes inside microglia pouches in the corpus callosum of juvenile WT and *cntnap2*^-/-^ mice. Histogram shows the total number of apoptotic oligodendrocytes, the proportion of Casp3^+^ oligodendrocytes and the proportion of them inside microglia pouches (n=4). Arrows point to apoptotic Olig2^+^ oligodendrocytes, some of the Casp-3^+^. Scale bar = 20 μm and 5 μm (higher magnifications). Statistical analysis was performed with 2-tailed unpaired Student t-test. *p<0.05, **p<0.01, ***p<0.001.

To identify the underlying molecular alterations, we performed bulk RNA-seq on MACS-isolated O4⁺ oligodendrocytes (encompassing premyelinating and myelinating stages) at P20 and P60. Principal component analysis revealed clear genotype separation at P20 (Fig. 2B), with 444 differentially expressed genes (DEGs; FDR-adjusted p < 0.05; no fold-change threshold applied) in *cntnap2*^-/-^ relative to controls (Fig. 2B). No significant DEGs were detected at P60 (Fig. S3A), consistent with phenotypic normalization in adults. KEGG pathway analysis (Fig. S4A) and gene set enrichment analysis (GSEA) identified two major dysregulated programmes at P20: upregulation of immune-related genes and downregulation of synaptic signalling pathways (Fig. 2C), neither of which persisted in adulthood (Fig. S3B). GSEA revealed significant enrichment of interferon-γ response (Fig. 2D), interferon-α response and TNFα/NF-κB signalling (Fig. S54B) in juvenile *cntnap2*^-/-^ oligodendrocytes. Key contributing genes included interferon-regulated transcription factors and effectors (*Irf7, Irf9, Ifih1, Ifit3, Ifit3b*) and MHC class I molecules (*H2-D1, H2-K1, H2-T23, B2m*). This transcriptional signature closely resembles the interferon-responsive oligodendrocyte state identified in multiple sclerosis lesions^38^, although here it occurs transiently during a critical developmental window rather than in the context of chronic demyelination. Confocal microscopy confirmed elevated expression of IFITM3, a canonical marker of interferon-responsive cells^38,39^, in *cntnap2*^-/-^ oligodendrocytes at P20 (Fig. 2E). This oligodendrocyte state has been associated with both antigen presentation capacity and endoplasmic reticulum stress^38^, raising the possibility that it actively impairs differentiation and myelination rather than solely reflecting a reparative response.

Since interferon-γ is primarily produced by microglia and infiltrating immune cells, we next examined microglial populations across the same developmental windows. At P20, *cntnap2*^-/-^ mice exhibited transient microgliosis, increased Iba1⁺ cell density in both S1 cortex and corpus callosum, that resolved by adulthood (Fig. 2F). Despite similar increases in density, microglial morphology differed markedly between regions. In S1 cortex, Sholl analysis revealed increased branching complexity and enlarged cell area (Fig. S5A), consistent with a hyper-ramified, surveillance-associated phenotype. In contrast, corpus callosum microglia displayed an amoeboid morphology with elevated iNOS expression (Fig. S5B), indicative of pro-inflammatory activation. This pro-inflammatory microglial state in the corpus callosum was associated with increased oligodendrocyte death. Apoptotic Olig2⁺ oligodendrocytes, identified by nuclear condensation, were more abundant at P20 (white arrows, Fig. 2G). Confocal imaging revealed two distinct populations of compromised oligodendrocytes within microglial pouches: Casp3⁺ cells with condensed nuclei, consistent with canonical apoptosis, and Casp3⁻ cells with intact nuclear morphology (orange arrows, Fig. 2G), raising the possibility of phagocytosis of viable oligodendrocytes^40^. Oligodendrocyte numbers in the corpus callosum showed a trend toward reduction at P20, with this deficit persisting significantly into adulthood (Fig. S5C). In S1 cortex, increased apoptotic oligodendrocyte death was also detected by Hoechst staining (Fig. S5D), although Casp3⁺ cells within microglial pouches were not observed, suggesting region-specific mechanisms of oligodendrocyte loss. Together, these data reveal transient developmental alterations in microglial state that coincide with the emergence of interferon-responsive oligodendrocytes and may contribute to the myelination deficits in *cntnap2*^-/-^ mice.

### Downregulation of GABAergic pathways in oligodendrocytes is associated with PV⁺ interneuron hypomyelination

In contrast to immune pathways, *cntnap2*^-/-^ oligodendrocytes exhibited marked downregulation of synaptic signalling gene clusters (Fig. S4A). Both GABAergic and glutamatergic pathways were significantly downregulated at P20 (Fig. 4A, B). Among GABAergic genes, reductions spanned multiple GABA-A receptor subunits (Gabra1–5, Gabrb1–3, Gabrg1–3), the GABA-B receptor subunit Gabbr2, and the GABA-synthesizing enzymes Gad1 and Gad2 (Fig. 3A). Glutamatergic signalling genes were also differentially expressed, including vesicular glutamate transporters (Slc17a7, Slc17a6), glutamate transporters (Slc1a2, Slc1a3), and ionotropic and metabotropic receptor subunits (Gria1, Grik1, Grin2d, Grm3) (Fig. 3B). However, whereas GABAergic receptor genes showed uniform and robust downregulation, glutamatergic receptor subunits displayed a mixed pattern of both up-and downregulation (Fig. 3C), pointing to a preferential disruption of GABAergic signalling in *cntnap2*^-/-^ oligodendrocytes.

**Figure 3.**
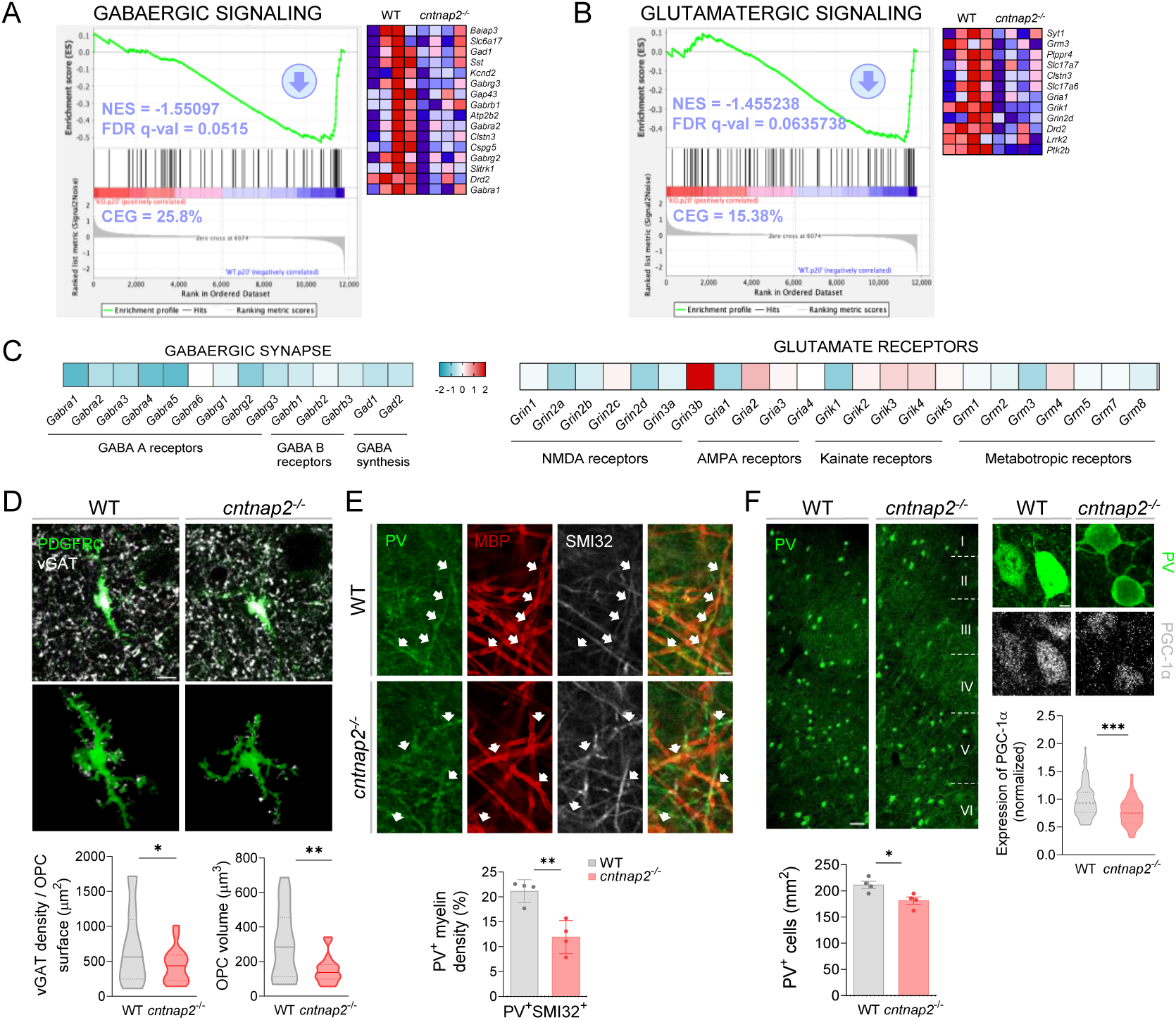
Downregulation of GABA signalling in oligodendrocytes leads to hypomyelination of PV^+^ interneurons. **A, B**) Enrichment plots (left) and heatmaps (right) of GABAergic and glutamatergic gene sets from hallmark analysis with corresponding NES, FDR q-value and Core Enrichment Genes (CEG). **C**) DEG of GABAergic and glutamatergic receptors between WT and *cntnap2*^-/-^ O4^+^ oligodendrocytes from juvenile mice. **D**) Confocal z stack images and 3D reconstructions of of vGAT^+^ presynaptic terminals (white) contacting oligodendrocyte precursor cells (OPCs, green). Scale bar= 5 μm. Below, vGAT^+^ presynaptic-OPC contacts and OPC volume in juvenile WT and *cntnap2*^-/-^ mice (n= 18-20 cells from 3 WT mice and 4 *cntnap2*^-/-^ mice). Scale bar= 5 μm. **E**) Confocal z stack images and quantification of myelin in PV^+^ interneurons axons (PV+, green; SMI32^+^, white) in S1 layer IV/V of juvenile *cntnap2*^-/-^ mice (n=4/4). Data was normalized to the density of PV^+^ SMI32^+^ axons. Scale bar= 3 μm. **F**) Left, confocal z stack images of PV+ interneurons and quantifications of their number in the total S1 of juvenile WT and *cntnap2*^-/-^ mice (n=4/4). Scale bar = 50 μm. Rigth, confocal z stack images of PGC-1a transcription factor and quantification in PV^+^ interneurons and quantifications in S1 layer V of juvenile WT and *cntnap2*^-/-^ mice (n= 50 cells from 4 mice per genotype). Scale bar = 5 μm. Data are represented as mean + SEM. Statistical analysis were performed with 2-tailed unpaired Student t-test. *p<0.05, **p<0.01

**Figure 4.**
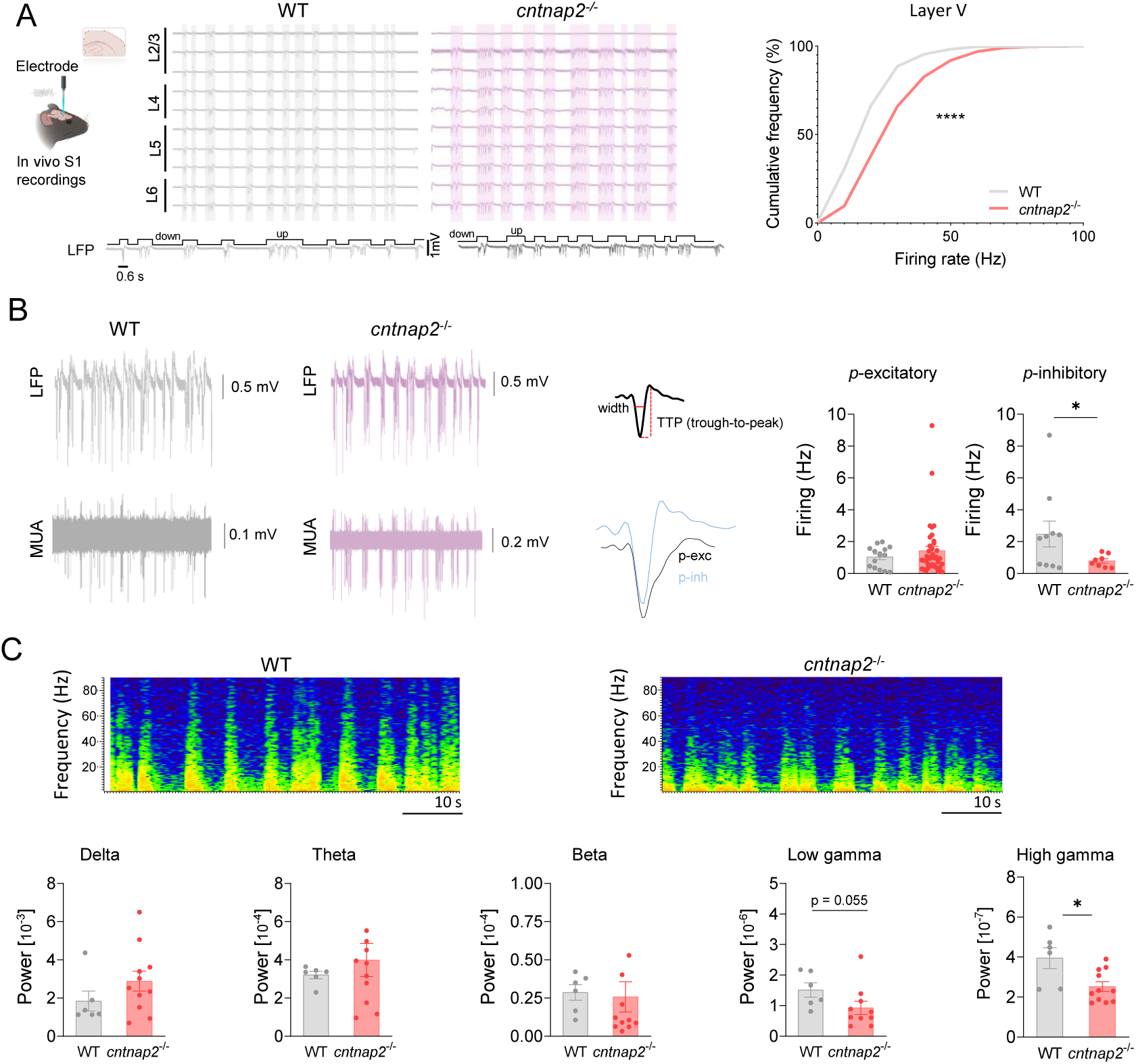
Impaired inhibitory activity in S1 of juvenile *cntnap2*^-/-^ mice. **A**) *Left*, schematic and LFP traces of the in vivo recordings in S1 of anesthetized mice in juvenile WT (gray) and *cntnap2*^-/-^ mice (pink). Gray and pink boxes indicate up-states. Bottom, raw LV traces in higher scale. *Right*, cumulative probability of spontaneous neuronal firing during up-states for juvenile WT and *cntnap2*^-/-^ mice (n= 6/11 for WT and *cntnap2*^-/-^ mice). **B**) Left, representative LFP and MUA with the respective spike units. Middle, representative half-width and through-to-peak-time (TTP) from the putative inhibitory and excitatory spike waveform. Right, firing rate of excitatory and inhibitory neurons during up-states (WT: n=15 excitatory neurons and n= 10 inhibitory neurons. *Cntnap2*^-/-^ mice: n= 39 excitatory neurons and n= 8 inhibitory neurons). **C**) LFP power spectra and analysis at different frequency bands: delta band (1-4 Hz), beta band (13-30 Hz), low gamma band (30-60 Hz) and high gamma band (60-100 Hz) in juvenile WT and *cntnap2*^-/-^ mice (n= 6/11). Data are represented as mean ± SEM. Statistical analysis were performed with Kolmogorov-Smirnov test (A) and unpaired Student t-test (B, C). *p<0.05, **p<0.01, ***p<0.001.

During early postnatal development, GABAergic communication between PV⁺ interneurons and OPCs is essential for activity-dependent myelination of inhibitory circuits. Deletion of GABA-A or GABA-B receptors in OPCs impairs PV⁺ interneuron myelination, leading to altered firing properties and deficits in cognition and social behaviour^32,33^. We therefore asked whether reduced GABAergic receptor expression in *cntnap2*^-/-^ oligodendrocytes impacts interneuron–OPC communication and PV⁺ interneuron myelination. To assess inhibitory input onto OPCs, we quantified vGAT⁺ presynaptic terminals apposed to three-dimensionally reconstructed PDGFRα⁺ OPC processes in S1 cortex (defined as being within a distance of 240 nm)^41^. The density of vGAT⁺ terminals contacting OPCs was significantly reduced in *cntnap2^-/-^* mice at P20 (Fig. 3D), indicating decreased GABAergic input. OPC volume was also significantly reduced in *cntnap2*^-/-^ mice, consistent with diminished inhibitory input to this population (Fig. 3D).

Among cortical interneurons, fast-spiking PV⁺ interneurons, predominantly located in layers IV and V, are the principal population to undergo myelination, which supports their high-frequency firing and precise temporal coordination^34^. We therefore examined whether PV⁺ interneuron myelination is selectively affected during development. Triple immunostaining for MBP, SMI32, and PV revealed a marked reduction in myelination of PV⁺ interneuron axons in layer V of S1 at P20 in *cntnap2*^-/-^ mice (Fig. 3E). In parallel, the number of PV⁺ interneurons in S1 was significantly reduced at P20 (Fig. 3F) but normalized in adults (Fig. S6). We next asked whether the intrinsic maturation programme of PV⁺ interneurons is also compromised. PGC-1α (encoded by Ppargc1a) is an activity-dependent transcriptional coactivator recently identified as a master regulator of PV⁺ interneuron terminal differentiation, governing the acquisition of fast-spiking properties, PV expression, and perineuronal nets^42^. Immunofluorescence analysis at P20 revealed that nuclear PGC-1α levels were significantly reduced in PV⁺ interneurons of juvenile *cntnap2^-/-^* mice compared with controls (Fig. 3F), indicating that developmental hypomyelination is accompanied by disruption of the transcriptional programme governing PV⁺ interneuron maturation.

Together, these findings demonstrate that *cntnap2*^-/-^ mice exhibit reduced GABAergic input onto OPCs, selective hypomyelination of PV⁺ interneurons, and impaired PV⁺ terminal differentiation, suggesting that these processes are temporally linked during a critical developmental window.

### PV⁺ interneuron hypofunction drives cortical hyperexcitability in juvenile *cntnap2^-/-^*mice

Loss of PV⁺ interneuron myelin induces synchronized epileptiform discharges through impaired PV-mediated inhibition^34^, and an elevated excitation–inhibition (E/I) ratio is a shared feature across genetic ASD models^6,7^. We therefore examined cortical excitability and inhibitory function in juvenile *cntnap2*^-/-^ mice by recording local field potentials (LFPs) from layer V of S1 in vivo. Spontaneous firing rates during up states were markedly increased in juvenile *cntnap2*^-/-^ mice (Fig. 4A), consistent with previous reports^6,7,25^ and indicative of enhanced cortical excitability. To identify the neuronal populations contributing to this phenotype, we performed spike sorting of individual single units and classified neurons as putative excitatory (p-exc) or putative inhibitory (p-inh) based on waveform half-width and trough-to-peak duration (Fig. 4B; 45). Firing rates of p-inh neurons were significantly reduced during up states in juvenile *cntnap2*^-/-^ mice (Fig. 4B), whereas p-exc firing rates showed a trend toward an increase, indicating an early developmental shift in the cortical E/I ratio.

Gamma oscillations depend on recurrent excitatory–inhibitory loops between pyramidal neurons and PV⁺ interneurons and are strongly phase-locked to PV⁺ interneuron spiking^8,43^. Given the link between PV⁺ interneuron myelination and gamma-band activity^34,44^, we assessed whether PV⁺ hypomyelination and hypofunction were associated with alterations in gamma oscillations. Spectral analysis of LFPs revealed a significant reduction in high-gamma power and a trend toward reduced low-gamma power (p = 0.058) in juvenile *cntnap2^-/-^* mice (Fig. 4C). Together, these findings link developmental myelination deficits to impaired inhibitory function and altered network dynamics in *cntnap2*^-/-^ mice.

### Clemastine rescues myelin deficits and restores E/I balance in layer V of somatosensory cortex in *cntnap2*^-/-^ mice

We next asked whether promoting oligodendrocyte differentiation during early development could reverse circuit abnormalities in *cntnap2*^-/-^ mice. Mice were treated with clemastine, a well-established promyelinating compound that promotes oligodendrocyte differentiation via muscarinic receptor 1^45^, for two weeks (P15–P29). Clemastine treatment promoted oligodendrocyte differentiation and significantly increased both the myelinated area of S1 and the number of nodes of Ranvier in layer V (Fig. 5A), confirming effective restoration of cortical myelination. In parallel, clemastine rescued the hypomyelination of PV⁺ interneuron axons observed in juvenile *cntnap2*^-/-^ mice (Fig. 5B). Notably, enhanced myelination was accompanied by a significant increase in PV⁺ interneuron numbers in S1 (Fig. 5C), suggesting that myelination supports PV⁺ interneuron survival and/or maturation during development. Consistent with this, clemastine restored PGC-1α expression in PV⁺ interneurons of *cntnap2*^-/-^ mice to wild-type levels (Fig. 5C). Given that PGC-1α is the master transcriptional regulator of PV⁺ interneuron terminal differentiation^42^, the rescue of its expression by a promyelinating intervention indicates that restoring oligodendrocyte differentiation is sufficient to reinstate not only myelin integrity but also the intrinsic transcriptional programme driving PV⁺ interneuron maturation.

**Figure 5.**
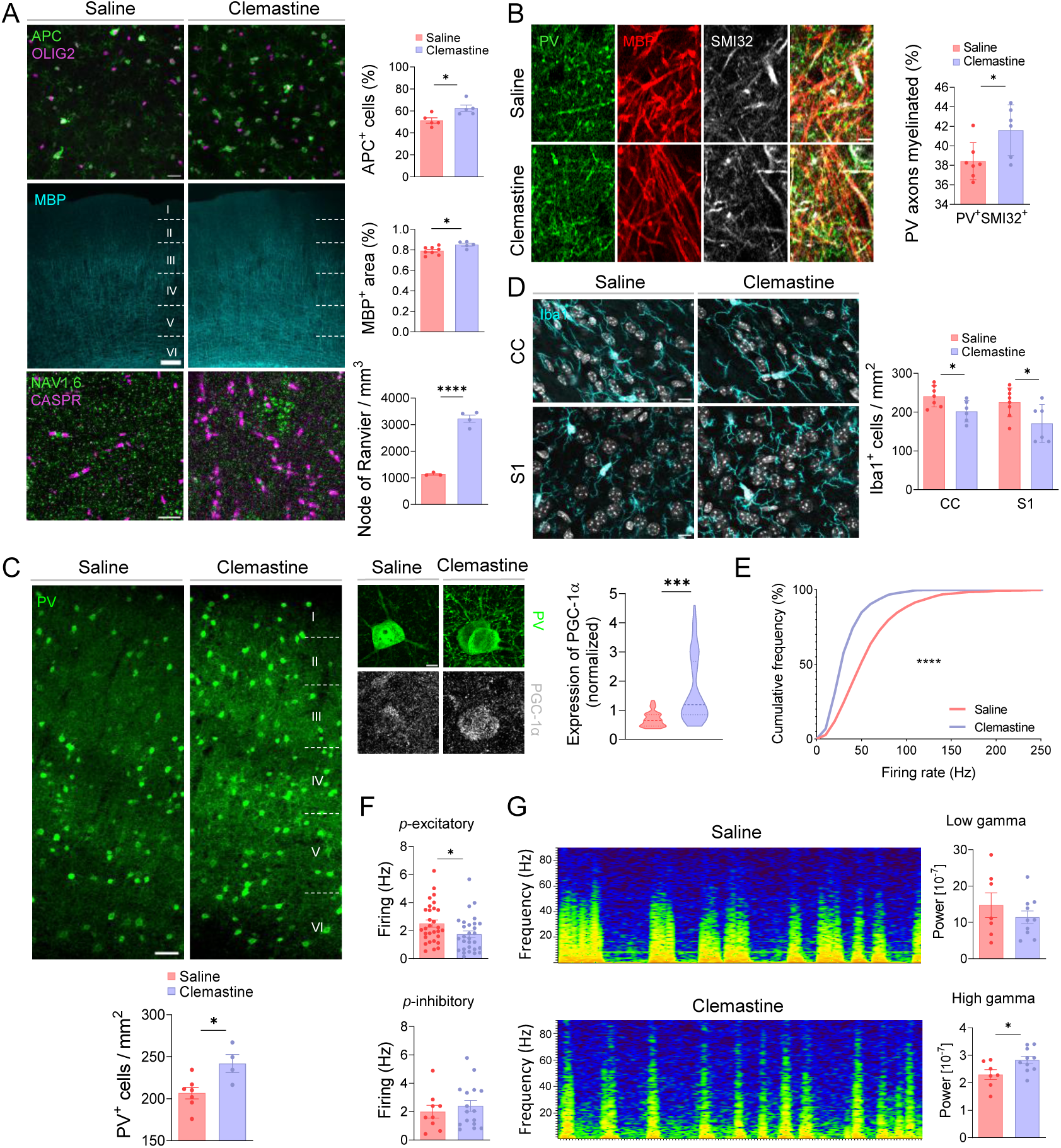
Clemastine treatment during development reverse myelin deficits and hyperexcitability of *cntnap2*^−/−^ mice. **A**) Confocal z stack images and quantifications of the proportion of mature APC^+^ oligodendrocytes (green) normalized to total Olig2^+^ oligodendrocytes (magenta) (top, n= 5/5), the S1 area myelinated (middle, n= 8/5) and the number of nodes of Ranvier (bottom, n= 3/4) in *cntnap2*^-/-^ mice treated with saline or clemastine. Scale bar = 25 μm (top), 50 μm (middle) and 5 μm (bottom). **B**) Confocal z stack images and quantification of myelin in PV^+^ interneurons axons (PV+, green; SMI32^+^, white) in the S1 layer V of *cntnap2*^-/-^ mice treated with saline or clemastine (n=7/6). Scale bar = 5 μm. **C**) Left, confocal z stack images and quantification of the number of PV^+^ interneurons in the S1 of *cntnap2*^-/-^ mice treated with saline or clemastine (n=7/6). Scale bar = 50 μm. Rigth, confocal z stack images and quantification of PGC-1α expression in layer V of S1 in *cntnap2*^-/-^ mice treated with saline or clemastine (n=70-75 cells from 7/6 mice). Scale bar = 5 μm. **D**) Confocal z stack images and quantifications of the number of microglia cells in corpus callosum and layer V of S1 in *cntnap2*^-/-^ mice treated with saline or clemastine (n =7/6). Scale bar = 10 μm. **E**) Cumulative probability of spontaneous neuronal firing during up-states for *cntnap2*^-/-^ mice treated with saline or clemastine (n= 7/10). **F**) Half-width and through-to-peak-time (TTP) from the averaged spike waveform for each cluster. Bottom, firing rate of excitatory and inhibitory neurons during up-states in *cntnap2*^-/-^ mice treated with saline or clemastine (Saline *cntnap2*^-/-^ mice: n= 31 excitatory neurons and 9 inhibitory neurons. Clemastine *cntnap2*^-/-^ mice: n= 28 excitatory neurons and 16 inhibitory neurons). **G**) LFP power spectra and analysis at different high frequency bands: low-gamma band (30-60 Hz) and high-gamma band (60-100 Hz) in juvenile *cntnap2*^-/-^ mice treated with saline or clemastine (n= 7/10). All data are represented as mean ± SEM. Statistical analysis were performed with Kolmogorov-Smirnov test (E) and two-tailed unpaired Student t-test (A,B C, D, F, G). *p<0.05, **p<0.01, ***p<0.001.

Beyond its effects on the oligodendroglial lineage, clemastine reduced microglial density in S1 of *cntnap2*^-/-^ mice (Fig. 5D), indicating attenuation of the transient developmental microgliosis. Similar effects on myelination and microglia were obtained with pranlukast (Fig. S7), an independent promyelinating compound acting through the orphan receptor GPR17^46^. Together, these findings demonstrate that promoting oligodendrocyte differentiation during a critical developmental window is sufficient to restore PV⁺ interneuron myelination, maturation, and number, while attenuating microglial activation.

To determine whether these cellular improvements translate into functional circuit rescue, we performed in vivo electrophysiological recordings in clemastine-treated *cntnap2*^-/-^ mice. Recordings from layer V of S1 revealed that clemastine significantly reduced overall cortical hyperexcitability (Fig. 5E). Spike-sorting analysis showed that clemastine decreased the firing rate of putative excitatory neurons compared with saline-treated *cntnap2*^-/-^ mice (Fig. 5F), whereas putative inhibitory neuron firing was not significantly altered (Fig. 5F). Although individual interneuron properties were not restored, the increased number of PV⁺ interneurons may provide sufficient cumulative inhibition to reduce excitatory firing. Accordingly, clemastine reversed deficits in high-gamma oscillation power (Fig. 5G). These results indicate that pharmacological enhancement of myelination ameliorates cortical hyperexcitability and partially normalizes E/I imbalance in *cntnap2*^-/-^ mice, though it is not sufficient to fully rescue local circuit dysfunction at the single-neuron level.

### Promoting myelination restores social deficits in *cntnap2*^-/-^ mice

We next asked whether promoting myelination during development could reverse behavioural alterations in *cntnap2^-/-^* mice. Mice were treated with clemastine or pranlukast during the same developmental window (P15–P29) and subsequently assessed in a comprehensive behavioural battery covering locomotor activity, anxiety-related behaviour, sensory responses, and social behaviour (Fig. 6A). Neither clemastine nor pranlukast altered locomotor activity in the open field test (Fig. S8A), indicating that promyelinating treatment does not rescue the hyperactive phenotype of *cntnap2^-/-^* mice. Similarly, neither compound affected centre–periphery exploration in the open field (Fig. S8A) or light–dark preference (Fig. S8B), confirming no effect on anxiety-related behaviour.

**Figure 6.**
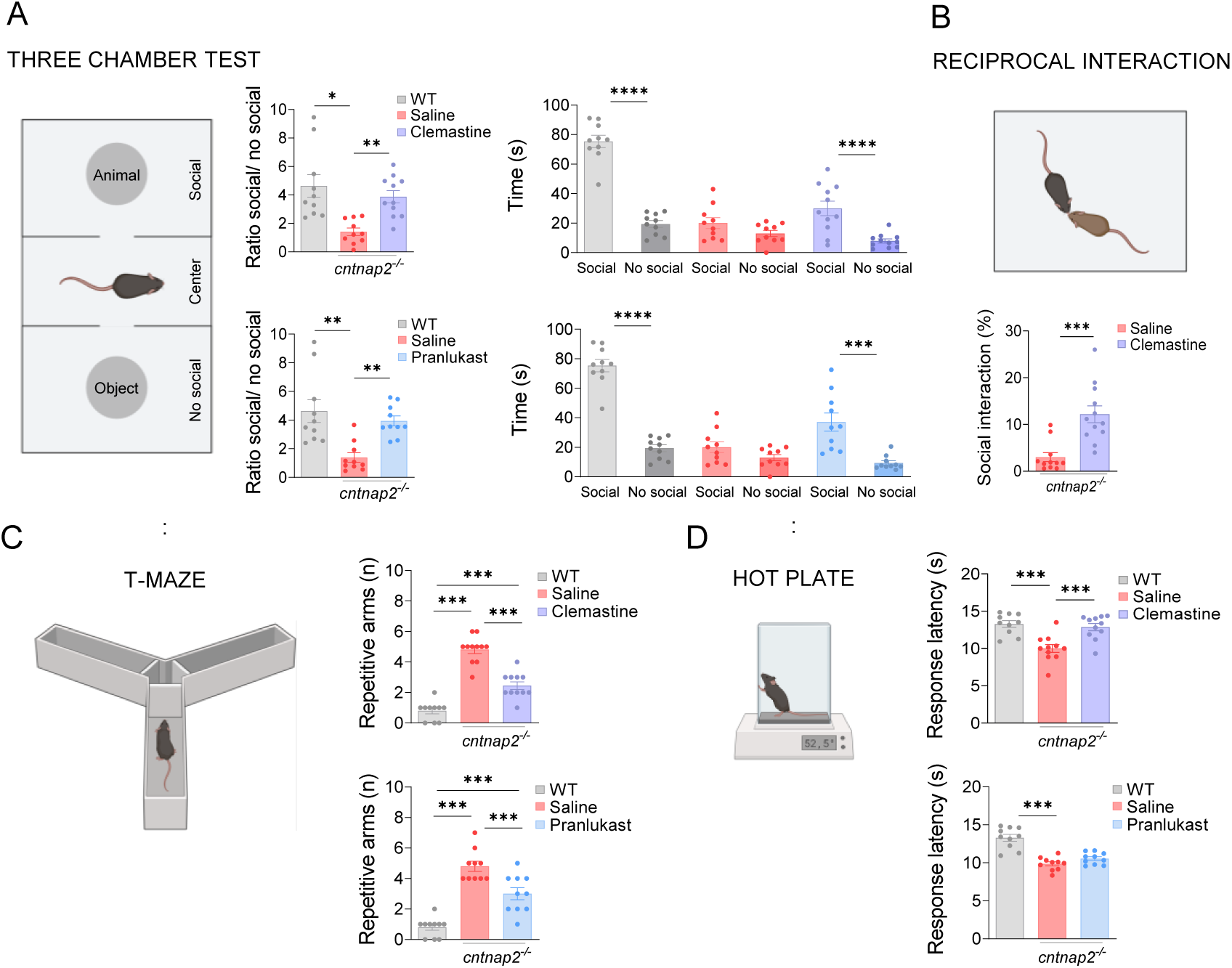
Clemastine and pranlukast treatment during development reverse the social deficits in *cntnap2*^−/−^ mice. **A**) Time spent exploring the social or non-social chamber and the ratio between them in WT (n=10) and *cntnap2*^-/-^ treated with saline, clemastine or pranlukast (n= 10/10/11 respectively). **B**) Time interacting with a mice matched in sex and age in WT mice ( n= 10) and in *cntnap2*^-/-^ mice treated with saline or clemastine (n= 10/10/11). **C**) Number of repetitions in the T-maze test in in WT (n= 9) and *cntnap2*^-/-^ treated with saline, clemastine or pranlukast (n= 10/10/11 respectively). **D**) Latency response in hot place test in WT mice (n= 10) and *cntnap2*^-/-^ mice treated with saline, clematine or pranlukast (n= 10/10/11 respectively). All data are represented as mean ± SEM. Statistical analysis were performed with ANOVA and Tukey’s multiple comparison post-hoc test (A, D), Kruskal-Wallis multiple comparison test and Dunn’s multiple comparison post-hoc test (C) and two-tailed non-paired t-Student (D). *p<0.05, **p<0.01, ***p<0.001.

In the three-chamber sociability test, untreated *cntnap2^-/-^* mice failed to show a preference for a social stimulus over an empty cup, confirming impaired sociability (Fig. 6B). Treatment with either clemastine or pranlukast restored this preference, as *cntnap2^-/-^* mice spent significantly more time interacting with the cup containing the stimulus mouse (Fig. 6B). This rescue was corroborated in the reciprocal social interaction test, in which clemastine significantly increased the total duration of social interactions between unfamiliar, genotype-and treatment-matched mice (Fig. 6C).

We next assessed repetitive behaviour using the spontaneous alternation T-maze test. Whereas control mice exhibited normal exploratory alternation between arms, *cntnap2^-/-^* mice displayed a marked increase in repetitive arm choices (Fig. 6D). Treatment with either clemastine or pranlukast significantly reduced the number of non-alternations across a 10-trial session (Fig. 7D), indicating attenuation of repetitive behaviour. Finally, sensory processing was evaluated using the hot plate assay. Clemastine, but not pranlukast, normalized the enhanced thermal sensitivity observed in *cntnap2^-/-^* mice (Fig. 6E).

Together, these results demonstrate that early postnatal promyelinating treatment rescues core ASD-relevant behavioural phenotypes such as social deficits and repetitive behaviour in *cntnap2^-/-^* mice, while hyperactivity and sensory alterations remain partially resistant to this intervention.

## DISCUSSION

In this study, we identify oligodendrocyte dysfunction as a key determinant of inhibitory circuit maturation and behavioural outcomes in the *cntnap2*^-/-^ mouse model of ASD. We show that transiently impaired OPC differentiation during early postnatal development leads to hypomyelination of PV⁺ interneurons, selective PV⁺ hypofunction, E/I imbalance, disrupted gamma oscillations, and social and repetitive behavioural deficits. Pharmacological promotion of myelination during this critical window reverses these circuit and behavioural abnormalities. These findings establish, for the first time in a genetically defined ASD model, a mechanistically complete and pharmacologically reversible chain linking oligodendrocyte dysfunction to inhibitory circuit maturation and autism-relevant behaviour.

Previous studies have reported reduced PV⁺ interneuron density, diminished inhibitory input onto pyramidal neurons, and altered gamma oscillations in *cntnap2*^-/-^ mice^15,43^, but the underlying mechanisms remained unclear. Our data identify developmental myelination as a critical upstream factor: PV⁺ interneuron hypofunction and reduced gamma oscillatory activity were both restored by pharmacological promotion of myelination. Notably, clemastine did not increase the firing rate of individual inhibitory neurons but instead expanded the PV⁺ interneuron population, likely by promoting maturation and PV expression. We propose that this population-level expansion, combined with restoration of myelin-dependent conduction properties, enhances the synchronization and temporal precision of inhibitory output onto pyramidal neurons — parameters that are more critical for gamma rhythm generation than the mean firing rate of individual units^8,34^. This interpretation aligns with computational models in which network synchrony, rather than firing rate per se, is the primary driver of gamma power^8^. Consistent with our findings, PV⁺ interneuron hypomyelination and impaired high-gamma oscillations have been independently reported in neuroligin-3 mutant mice^25^, and cuprizone-induced demyelination experiments have demonstrated that PV⁺ axon myelination is required to stabilize fast inhibitory transmission and regulate gamma oscillations^34,44^. Together, these convergent lines of evidence position myelin as a critical modulator of inhibitory circuit function in the context of neurodevelopmental disorders. Although myelin deficits in *cntnap2*^-/-^ mice were not restricted to PV⁺ interneurons — reduced myelination of excitatory callosal projections was also observed — our electrophysiological data indicate that PV⁺ interneurons are disproportionately affected. Pyramidal neurons exhibited increased firing rates, likely reflecting reduced inhibitory input rather than a direct consequence of their own hypomyelination^7^. The selective vulnerability of PV⁺ interneurons may be explained by at least three non-mutually exclusive mechanisms. First, the exceptionally high energetic demands of fast-spiking activity render PV⁺ interneurons particularly sensitive to the metabolic support provided by myelinating oligodendrocytes^47,48^. Second, PV⁺ interneuron myelination occurs during a narrow early postnatal window that coincides precisely with the period of hypomyelination identified here, making these cells disproportionately vulnerable to disruptions at this developmental stage^13,32,33^. Third, the bidirectional GABAergic communication between PV⁺ interneurons and OPCs — which we show to be reduced in *cntnap2*^-/-^ mice — has no functional equivalent for pyramidal neurons, rendering PV⁺ interneuron myelination uniquely dependent on this activity-dependent signalling axis. Consistent with this framework, hypomyelination secondary to loss of ErbB receptor signalling in oligodendrocytes leads to reduced PV expression and impaired inhibitory synaptic function without altering excitatory connectivity^49^.

A central finding of this study is that developmental hypomyelination in *cntnap2*^-/-^ mice is accompanied by reduced PGC-1α expression in PV⁺ interneurons, and that promoting myelination with clemastine is sufficient to restore it. PGC-1α has recently been identified as an activity-dependent master regulator of PV⁺ interneuron terminal differentiation, orchestrating the expression of PV, perineuronal nets, voltage-gated potassium channels (Kv3.1, Kv1.1), and mitochondrial biogenesis genes through a transcriptional complex with ERRγ and Mef2c^42^. The reduction in PGC-1α at P20 therefore indicates a broader failure of the intrinsic maturation programme of PV⁺ interneurons, extending beyond the loss of PV as a single marker. The rescue of PGC-1α expression by a promyelinating intervention establishes a functional link between oligodendrocyte-mediated myelination and the activation of PGC-1α-dependent terminal differentiation. Since PGC-1α is upregulated in response to increasing neuronal activity during the second postnatal week^42^, one possibility is that hypomyelination impairs PV⁺ interneuron firing properties, reducing the activity-dependent signals required to drive PGC-1α expression, creating a self-reinforcing loop in which deficient myelination prevents the maturation programme that would in turn support further myelination and fast-spiking function. Alternatively, myelin-derived metabolic or trophic signals may directly contribute to PGC-1α induction, a possibility that warrants future investigation.

Our transcriptomic analysis reveal that the most pronounced molecular alterations in oligodendrocytes occur during the early postnatal period, with substantial normalization in adulthood. This developmental specificity aligns with the emerging view that oligodendrocytes actively shape the maturation and stabilization of cortical circuits, and that developmental myelination acts as a functional brake defining the closure of critical periods^50,51^. Crucially, despite molecular recovery, social deficits persist in adult *cntnap2*^-/-^ mice, indicating that disruptions during this critical window exert enduring consequences on brain function and behaviour that cannot be reversed by delayed normalization of myelin parameters^52^. This observation is consistent with previous findings showing that transient developmental perturbations to PV⁺ interneurons — including excess PV⁺ cell numbers^52^ or disrupted PV–OPC communication via GABA-A γ2 subunit deletion^33^ — lead to persistent behavioural deficits even when cellular parameters later normalize. Together, these data underscore the irreversibility of circuit-level consequences when myelination is disrupted during a critical developmental period, and highlight the importance of early intervention.

An important unresolved question concerns the cellular origin of the myelination deficits. The preserved capacity of *cntnap2*^-/-^ oligodendrocytes to differentiate and myelinate nanofibers in vitro indicates that the impairment arises primarily from extrinsic cues within the CNS parenchyma rather than from cell-autonomous dysfunction, as described for certain other ASD-linked genes ^22,23,25,53^. Since *cntnap2* is predominantly expressed in axons with minimal expression in oligodendrocytes, disrupted axon–oligodendrocyte signalling and altered microenvironmental cues are the most compelling explanations. Two non-mutually exclusive mechanisms emerge from our data. First, the upregulation of interferon signalling pathways in *cntnap2*^-/-^ oligodendrocytes, together with transient microgliosis and increased oligodendrocyte phagocytosis, suggests that altered neuroimmune signalling contributes critically to oligodendrocyte dysfunction during development. Microglia are essential regulators of developmental myelination^54–56^ and refine oligodendrocyte populations through a pruning process analogous to synaptic refinement^57,58^. The engulfment of both apoptotic and apparently viable oligodendrocytes observed in the corpus callosum suggests that pathological microglial over-pruning may directly reduce the pool of myelinating cells during the critical window. Notably, the resolution of microgliosis by clemastine and pranlukast — two compounds that act directly on oligodendrocyte differentiation rather than on microglial targets — suggests that the microglial activation is secondary to the myelination deficit rather than its primary cause, consistent with a reactive response to the disrupted oligodendroglial environment. Second, the reduced GABAergic synaptic input from interneurons onto OPCs, combined with downregulation of GABA receptor expression in *cntnap2*^-/-^ oligodendrocytes, would impair activity-dependent myelination of PV⁺ interneurons, creating a feed-forward loop in which reduced inhibitory output further diminishes the myelination signal to OPCs. Notably, GABA-responsive microglia have been shown to selectively sculpt inhibitory circuits during development^59^, providing a potential molecular link between these two mechanisms and suggesting that disrupted GABAergic signalling may simultaneously impair both oligodendrocyte-directed and microglia-directed refinement of inhibitory circuits.

The rescue of social deficits and behavioural inflexibility by clemastine and pranlukast — two mechanistically distinct promyelinating compounds — provides strong evidence that the behavioural phenotype is causally linked to myelination rather than to off-target pharmacological effects. However, neither compound rescued hyperactivity, and the sensory phenotype (enhanced thermal sensitivity) was normalized by clemastine but not pranlukast — a dissociation that may reflect the distinct pharmacological profiles of these compounds beyond their shared promyelinating activity, as clemastine also acts as an antihistamine (H1 receptor antagonist), which could independently modulate nociceptive processing. These findings indicate that while myelination is sufficient to rescue social and cognitive phenotypes, other ASD-related features likely depend on additional circuit mechanisms that are not exclusively regulated by oligodendrocyte function.

Together, our findings identify disrupted developmental myelination as a critical mechanism linking immune signalling, oligodendrocyte dysfunction, and inhibitory circuit maturation to autism-relevant behavioural phenotypes. We propose that transient alterations in the CNS microenvironment during early postnatal development — encompassing neuroimmune activation, disrupted interneuron–OPC communication, and impaired oligodendrocyte differentiation — selectively compromise PV⁺ interneuron maturation and function, destabilizing E/I balance and derailing cortical circuit maturation during a critical period whose consequences persist into adulthood. The reversibility of these deficits through early promyelinating intervention highlights developmental myelination as a modifiable process and a compelling therapeutic target. More broadly, these results underscore the importance of glial–neuronal interactions during critical periods in shaping functional brain circuits, and establish that early disruptions to myelin-dependent circuit refinement have lasting consequences for neurodevelopmental disorders.

## MATERIALS AND METHODS

### Animals

Experiments were performed on C57/BL6 wildtype and *cntnap*2^-/-^ mice expressing a tau-LacZ fusion protein under the endogenous *cntnap2* promoter. The experimental procedures were approved by the Research Ethics and Animal Welfare Committee of the University of Basque Country (UPV/EHU) and are in accordance with the European Communities Council Directive (2010/63/UE) on the protection of animals used for scientific purposes. Mice were kept on a 12 h light/dark cycle with constant ambient temperature (22 ± 1 °C) and humidity, and water and food available ad libitum. Experiments were performed during the light phase of the cycle. Clemastine (50 mg/kg, Merck) and pranlukast (0.5 mg/kg, GenoChem) were administered daily intraperitoneally.

### Oligodendrocytes cultures

Primary oligodendrocyte cultures were obtained as previously^60^. Briefly, optic nerves were extracted from P11-12 rats. After enzymatic and mechanical digestion, cells were filtered and seeded at 10,000 cells/well on 1 μg/ml poly-D-lysine-coated (PDL) coatings (Sigma-Aldrich). Oligodendrocytes were maintained in SATO differentiation media^60^. For myelination assays, cells were seeded on coverslips with polycaprolactone nanofibers (Sigma-Aldrich) at a 20,000 cells/well density and CNTF and NT-3 were omitted from the medium. Cell differentiation and myelination in vitro was quantified in 4 different cultures performed in triplicate. In each experiment at 10 cells per field and 3 different fields were quantified per experiment.

### Immunohistochemistry

Mice were transcardially perfused with 4% p-formaldehyde (PFA) in 0.1 M sodium phosphate buffer (PBS) (pH 7.4) and post-fixated in 4% PFA overnight. Tissue was cut using a Microm HM650V vibratome (40 μm). After blocking with 4% normal goat serum in PBS plus 0.1% Triton X-100 (blocking solution) for 1 h at room temperature, free-floating sections were incubated with primary antibodies overnight at 4°C in blocking solution. The following antibodies were used: anti-APC (1: 500, #OP80 Sigma), anti-Olig2 (1:1000, #MABN50 Sigma), anti-caspr (1:500, #MABN69 Sigma), anti-MBP (1:1000, #808401 Biolegend), anti-Nav1.6 (1:500, #ASC-009 Alomone), anti-PDGFRα (1:250, #sc-398206 SantaCruz Biotechnology), anti-Iba1 (1:500, #019-19741 Wako), anti-BCAS1 (1:500, #445 004 Synaptic Systems), anti-cleaved caspase 3 (1:400, #9661 Cell Signaling), anti-iNOS (1:250, #610432 BD Biosciences), anti-PV (1:500, #NB120-11427 Synaptic systems), anti-PGC-1α (1:1000, #sc-518025 SantaCruz Biotechnology), anti-SMI32 (1:1000, #801701 Covance) and anti-IFITM3 (1:300, #11714-1-AP Proteintech). Tissue sections were washed three times with PBS and incubated with the appropriate secondary antibodies: anti-rat, anti-rabbit, anti-guinea pig, and anti-mouse IgG AlexaFluor-488,-594, and-647 secondary antibodies (1:250, Invitrogen) for 1 h at room temperature. Cell nuclei were stained using DAPI (4 μg/ml; Sigma-Aldrich). For in vitro immunocytochemistry, cells were fixed with 4% PFA for 20 min. After blocking with 4% normal goat serum in PBS plus 0.1% Triton X-100 (blocking solution) for 30 min, cells were incubated with primary antibodies in blocking solution for 2h at room temperature, washed and subsequently incubated with secondary antibodies for 1h at room temperature.

### Image acquisition and quantification

Images were acquired using identical acquisition settings for all samples within each experimental group on Zeiss LSM800, Zeiss LSM880 Airyscan, and Leica TCS SP8 confocal microscopes. Image analysis was performed using custom scripts and semi-automated workflows implemented in ImageJ and MATLAB. For preprocessing and generation of binary masks, the following FIJI functions were applied in batch-processing mode: *Subtract Background, Filter, Threshold, and Object Counter*. Myelin parameters and cell counts were quantified in 3-4 slices per mice. To quantify myelin coverage, we performed a three-dimensional analysis using the *Object Counter* and *3D Manager* plugins to segment myelin structures^66^, followed by *Skeletonize* and *Analyze Skeleton (3D)* functions to determine the total length of myelinated axons. Myelin was measured within a volume of 0.5-0.7 mm^3^ in S1 cortex and data was normalized to mm^3^. Orientation of myelin fiber was quantified using the *OrientationJ* Fiji plugin (https://github.com/Biomedical-Imaging-Group/OrientationJ). Myelination of PV axons was assessed by triple immunofluorescence labelling using antibodies against MBP, SMI32, and PV to label myelin, axons, and PV⁺ interneurons, respectively. Myelination of PV⁺ axons were quantified on 63X z-stack images using the BIOP JACoP plugin for ImageJ and normalized to the PV axonal area. The number and length of nodes of Ranvier were quantified from maximum-intensity projections of high-magnification (63×) z-stack images. A node was manually measured only when the node and both flanking paranodes were fully delineated using immunostaining for Caspr and Nav1.6. At least 25 nodes were quantified per mouse. The relative proportions of oligodendrocyte progenitor cells (OPCs) and mature myelinating oligodendrocytes were determined by triple immunofluorescence labelling using antibodies against PDGFRα, APC, and Olig2 to identify OPCs, mature oligodendrocytes, and total oligodendroglial lineage cells, respectively. Quantification was performed on 40× z-stack images. Oligodendroglial cell death was assessed by immunostaining for cleaved caspase-3 (Casp-3) combined with Hoechst nuclear staining to detect chromatin condensation, and values were normalized to the analyzed area or the total Olig2^+^ oligodendroglial cells. Microglial cell numbers were determined by immunostaining with anti-Iba1 and Sholl analysis was performed on high-magnification (63×) z-stack images. Individual microglial cells were segmented based on Iba-1 immunostaining, and binarized images were skeletonized to delineate cellular processes. Morphological complexity was defined as the number of intersections between microglial processes and concentric circles centered on the soma and spaced 5 µm apart. Analyses were performed on at least 15 cells per mouse. For iNOS expression analysis, a mask generated from the Iba1 channel was applied to high-magnification (63×) images acquired using a Leica TCS SP8 super-resolution microscope, ensuring that iNOS labeling was quantified exclusively within microglial cells. Analyses were performed on at least 15 cells per mice. To investigate communication between PDGFRα⁺ OPCs and interneurons, we quantified the number of vGAT⁺ presynaptic GABAergic terminals in direct apposition to the OPC surface (distance <240 nm). Confocal image stacks were first deconvolved using Huygens Professional software, after which PDGFRα⁺ cells were segmented using the *Trainable Weka Segmentation* plugin in FIJI^61^. Individual cells were further refined with thresholding and segmented in three dimensions with the *3D Object Counter* plugin to enable quantitative analysis. vGAT⁺ presynaptic terminals were automatically segmented using the FIJI 3D suite^62^. Subsequently, the *Selection > Make Band* tool and Boolean operations applied to PDGFRα and vGAT binary masks were used to automatically determine the number of contacting synapses in three dimensions. The analysis was conducted on a total of 20 cells from four different mice per experimental group. The expression of PGC-1a was assessed at single cell level and normalized to PV expression within each cell to account for variability in cell size. All quantifications were performed blinded to the experimental conditions to avoid bias in the analysis.

### Transmission electron microscopy

For the analysis of myelination trough transmission electron microscopy (TEM), mice were perfused with 4% PFA and 4 % glutaraldehyde in 0.1 M PB for 15 min. Brains were collected and post-fixed overnight at 4°C in the same solution. Subsequently, coronal 50 μm-thick brain sections were cut using a Microm HM650V vibrating blade microtome (Thermo Fisher, SGiker, EHU). Tissue sections were further fixed in 1% osmium tetroxide solution, dehydrated in increasing concentrations of ethanol and embedded in epoxy resin (Sigma Aldrich) to generate capsules containing the corpus callosum and the somatosensory cortex region. Semithin (1 μm-thick) and ultrathin (55 nm-thick) sections were obtained using an ultramicrotome RMC Boeckeler (SGIker, UPV/EHU). Then, ultrathin sections were contrasted by incubation in 2 % uranyl acetate solution for 10 min at RT and 0.2% lead citrate solution for 5 min at RT, to visualize the myelin structure. TEM images were obtained using the Zeiss EM900 electron microscope (SGiker, EHU). For the analysis of myelin sheath thickness, 8,000X magnification was used and at least 100 axons were analysed per animal. Outer fiber area, inner fiber area and axon area were measured and converted to diameter values to calculate g-ratios (inner fiber diameter/outer fiber diameter). In addition, inner tongue, and periaxonal space were quantified semiautomatically in 8,000X magnification images using MyelTracer^63^.

### Transcriptomic analysis

Oligodendrocytes were isolated from postnatal day (P) 20 and P60 wild-type (WT) and *cntnap2⁻/⁻* mice using magnetic-activated cell sorting (MACS) with O4-conjugated beads (Miltenyi Biotec). Animals were anesthetized with isoflurane and sacrificed by decapitation. Brains were rapidly removed and placed in Hibernate A medium (Gibco). The cerebral cortex was dissected and cut into ∼1 mm³ fragments. Tissue digestion was performed with a solution containing papain (165 U/ml) and DNase I (0.04 mg/ml) in Hibernate A for 30 minutes at 37°C. After tissue digestion, papain was removed by centrifugation in Hanks Buffered Salt Solution for 5 minutes at 300 x g (Gibco). The tissue was then triturated in Hibernate A supplemented with 2% B27 (Gibco) and 2 mM sodium pyruvate (Gibco). Four rounds of trituration were performed, using first a 5 mL pipette, polished glass Pasteur pipettes of decreasing diameter and a 1 mL pipette. In each round, the supernatant was kept and the remaining tissue was processed using fresh trituration solution. The resulting cell suspension was filtered through a 70 μm cell strainer (VWR) and centrifuged for 20 minutes at 800 × g in 22.5% Percoll solution (GE Healthcare) prepared in DMEM GlutaMAX (Gibco). The resulting pellet was washed with HBSS and resuspended in 10 mL of MACS solution containing 2 mM sodium pyruvate, 2% B27, 2 mM EDTA, 0.5% BSA and 10 μg/ml insulin in Hibernate-A, and incubated for 2 hours in petri dishes to remove microglia. The suspension was collected from the plates and centrifuged for 5 minutes at 300 x g. Next, cells were incubated in 100 μL MACS solution with anti-O4 antibody (1:10) (Milteny Biotech) for 15 minutes on ice. Excess antibody was removed by washing in HBSS and the cells were magnetically sorted following the manufacturer’s instructions (Milteny Biotech).

Total RNA was extracted from oligodendrocytes using the NZY Total RNA Isolation Kit (NZYTech), following the manufacturer’s protocol. RNA quantity and quality were assessed using the Qubit RNA HS Assay Kit (Thermo Fisher) and Agilent RNA 6000 Nano Chips (Agilent Technologies), respectively. RNA sequencing libraries were prepared using the TruSeq® Stranded mRNA Library Prep Kit and TruSeq RNA CD Index Plate (Illumina). From 400 ng of total RNA, mRNA was purified, fragmented, and primed for cDNA synthesis using Illumina and Thermo Fisher reagents. Libraries were then A-tailed, ligated to adaptors, and enriched by PCR. Library quality and concentration were evaluated using the Qubit dsDNA HS Assay Kit (Thermo Fisher Scientific) and visualized on an Agilent 2100 Bioanalyzer using the High Sensitivity DNA Kit (Agilent Technologies). Sequencing reads (FASTQ files) were aligned to the *Mus musculus* reference genome (“rn6”) using STAR version 2.7.10b. Read counts for annotated features were obtained from processed BAM files using Subread’s FeatureCounts v2.0.3. Differential expression analysis was performed with the DESeq2 R package (version 1.44.0). Gene set enrichment analysis (GSEA) was conducted using curated gene sets with 1,000 gene set permutations to assess significance, and genes were ranked using the signal-to-noise metric, according to the Molecular Signatures Database (MSigDB, https://www.gsea-msigdb.org/gsea/msigdb/index.jsp).

### Compound action potentials recordings

Mice were anesthetized and decapitated and the brains were rapidly removed and placed in ice-cold (4°C) cutting artificial cerebrospinal fluid (ACSF) containing (in mM): 215 Sucrose, 2.5 KCl, 26 NaHCO3, 1.6 NaH2PO4, 20 Glucose, 1 CaCl2, 4 MgCl2, and 4 MgSO4 bubbled with a mixture of 95% O2/5% CO2. Coronal slices (400-μm thick) were prepared using a Leica VT1200S vibratome and transferred to warmed (∼36 ° C) normal ACSF (nACSF) containing (in mM): 124 NaCl, 2.5 KCl, 10 glucose, 25 NaHCO3, 1.25 NaH2PO4, 2.5 CaCl2, and 1.3 MgCl2 for a 45-minute recovery. Compound action potentials (CAPs) were evoked by electrical stimulation of the corpus callosum with a bipolar electrode (CE2C55, FHC) and recorded with a pulled borosilicate glass pipette electrode (1.6 MΩ resistance) filled with NaCl 3 M, positioned in the contralateral corpus callosum (1000 μm distance from the stimulation site. Stimulation intensities ranged from 30 to 3000 μA (100 μs pulses, Master-8, AMPI). Response latency values for N1 and N2 fibers were used to calculate conduction velocity. At least three responses were averaged for each measurement and 3-4 slices were recorded per mouse. Peak amplitudes and latencies were calculated using custom routines in pCLAMP 10.0 (Molecular Devices).

### In vivo electrophysiology recordings and analysis

In vivo electrophysiology was be performed using multichannel electrode recording arrays (Neuronexus, 16 channels, horizontal custom-made array), as previously described^64,65^. Briefly, mice were anaesthetized and fixed in a stereotaxic with body temperature kept at 36.5°C using a homoeothermic blanket. Then, a 2×2-mm craniotomy was carried out over the right S1 (ML 1.3 to 1.8 mm, AP-0.3 to −0.8 mm) to lower the vertical array to record LFPs and APs across layers with a OmniPlex System (Plexon Inc, Dallas TX) and analyzed with Spike v.2, Matlab or IgorPro. Spontaneous activity was obtained in three separate recordings of four minutes each. Recordings were converted into digital data at a 40 kHz sampling rate, with 16–bit quantization by an OmniPlex System controlled by OmniPlex Server and PlexControl software (Plexon Inc, Texas, USA). Neural signals were processed offline using Spike2 software (v7, Cambridge Electronic Design, Cambridge, UK), and separated into two components: local field potentials (LFPs), representing low-frequency unfiltered signals, and multiunit activity (MUA), derived from high-frequency content filtered using a finite impulse response (FIR) band-pass filter (0.35–3 kHz, with a 0.12 kHz gap). Spike detection in the MUA signal was performed by applying a threshold of 3.5 standard deviations above the baseline voltage during down-states (>200 ms). The firing rate during up-states from the most active channel within layer 5 (600–850 μm) was calculated using the semi-automatic built-in script Bursts.s2s running on Spike2 v7.12. Spike sorting was performed on the MUA signal using the Cluster of Measurements algorithm in Spike2 v7. Spikes were extracted within a 7-ms window (−2 to +5 ms before and after the spike negative peak, respectively). Units were clustered based on two waveform features: trough-to-peak time (TTP, ms) and half peak height (Half-width, ms). Clusters with a trough-to-peak time <0.7 ms and half-width <0.3 ms were classified as putative inhibitory. Classification was supervised and manually corrected when necessary; spikes meeting only one inhibitory criterion were manually reassigned. Power spectrum analysis was performed in MATLAB R2024b (MathWorks, Natick, MA) using custom-written scripts. LFPs signals were analysed to quantify the distribution of oscillatory power across canonical frequency bands. Signals were first downsampled and bandpass filtered (1–100 Hz) to remove slow drifts and high-frequency noise. Power spectral density (PSD) was computed using Welch’s method with 2-s Hamming windows and 50% overlap. Mean power values were extracted for the delta (1–4 Hz), theta (4–12 Hz), beta (12–30 Hz), low gamma (30–60 Hz), and high gamma (60–100 Hz) frequency bands. For each recording, band-specific power was normalized to the total spectral power (1–100 Hz) to enable comparison across animals and experimental conditions. The theta/high-gamma ratio was computed as an index of the balance between low-frequency rhythmic synchronization and high-frequency cortical activation. LFP signals were filtered in the theta (4–12 Hz) and high-gamma (60–100 Hz) bands, and PSDs were estimated using the same Welch parameters described above. For each recording, the normalized mean power in the theta band was divided by the normalized mean power in the high-gamma band. Resulting ratios were averaged across recording sites for each animal and treatment.

### Behaviour analysis

Animals were treated intraperitoneally (i.p.) for 15 consecutive days during the critical developmental window (P15–P29) with two promyelinating drugs, clemastine and (50 mg/kg, Merck) and pranlukast (0.5 mg/kg, GenoChem). Following treatment, a comprehensive battery of behavioral tests was performed to assess potential effects on anxiety, locomotor activity, sensory response, social interaction, and cognitive flexibility. Open Field Test: Mice were placed in a clear Plexiglas arena (27.5 × 27.5 cm) for 10 minutes of habituation. After habituation, locomotor activity was recorded for 10 minutes, and general activity parameters—including total distance travelled, velocity, and movement patterns—were quantified. Reciprocal Social Interaction Test: Mice were placed in a cage previously habituated for 10 minutes alongside an unfamiliar, age-, sex-, and genotype-matched conspecific. Both mice in each pair received the same treatment (saline or clemastine). The total time engaged in social interactions—including nose-to-nose sniffing, nose-to-anus sniffing, following, or crawling on/under the partner—was measured for the pair. Three-Chamber Test: Following 10 minutes of habituation, a test mouse was placed in the central chamber of a clear Plexiglas apparatus divided into three interconnected chambers. The mouse was given the choice to interact with either an empty wire cup (located in one side chamber) or a similar cup containing an unfamiliar WT mouse (matched for age and sex) in the opposite chamber. Time spent interacting with each cup was recorded. T-Maze Spontaneous Alternation: Mice were placed at the base of a T-maze and allowed to explore either the right or left arm for ten consecutive trials. Entry was registered once all four paws entered an arm, at which point the gate to that arm was closed, and the animal was allowed to explore the arm for 5 seconds before returning to the start. Hot Plate Test: Mice were placed on a heated surface (52.5°C), and latency to paw withdrawal was measured, with a maximum cutoff of 15 seconds to prevent injury. All behaviours were recorded and analysed offline with Ethovision XT software (Noldus). No sex-specific differences in behaviour were detected; therefore, data from male and female mice were pooled, with comparable numbers of each sex included across all behavioural assays.

## Statistical Analyses

Data distribution normality was assessed using the Shapiro-Wilks test, and homogeneity of variances was evaluated with Levene’s test. Any outliers identified were removed prior to statistical analysis. All results are presented as mean ± SEM, except for violin plots, which depict the median and interquartile range. Statistical analyses were conducted using GraphPad Prism (version 9.0; GraphPad Software). Comparisons between two groups were performed using paired two-tailed Student’s t-tests, unless otherwise indicated. Comparisons between more than two groups were performed with ANOVA, unless otherwise indicated, and Turkey’s post-hoc test. Statistical significance was defined as p < 0.05.

## Acknowledgments

The authors thank the Animal Service and the Analytical and High-Resolution Facility of the UPV/EHU SGIker services at the University of the Basque Country (UPV/EHU), the Molecular Biology and Imaging Facilities at the Achucarro Basque Center for Neuroscience, and the Transcriptomics Service at CIC Biogune (Bizkaia, Spain).

## Author contributions

Conceptualization: MD

Methodology: MV, SC, LB-C, MVS-G, FNS, OF, OP, JRM

Investigation: MV, IU, AS-C, CP-M, AR, PM, YR, OP, JRM

Visualization: MV, IU, JMR, MD

Funding acquisition: MD, JMR

Project administration: MD

Supervision: MD, JMR

Writing – original draft: MD

Writing – review & editing: MD, FNS, JMR

