## Supplementary Figures for "Oligodendroglial dysfunction drives parvalbumin interneuron hypomyelination, circuit dysfunction and social deficits in an ASD model"

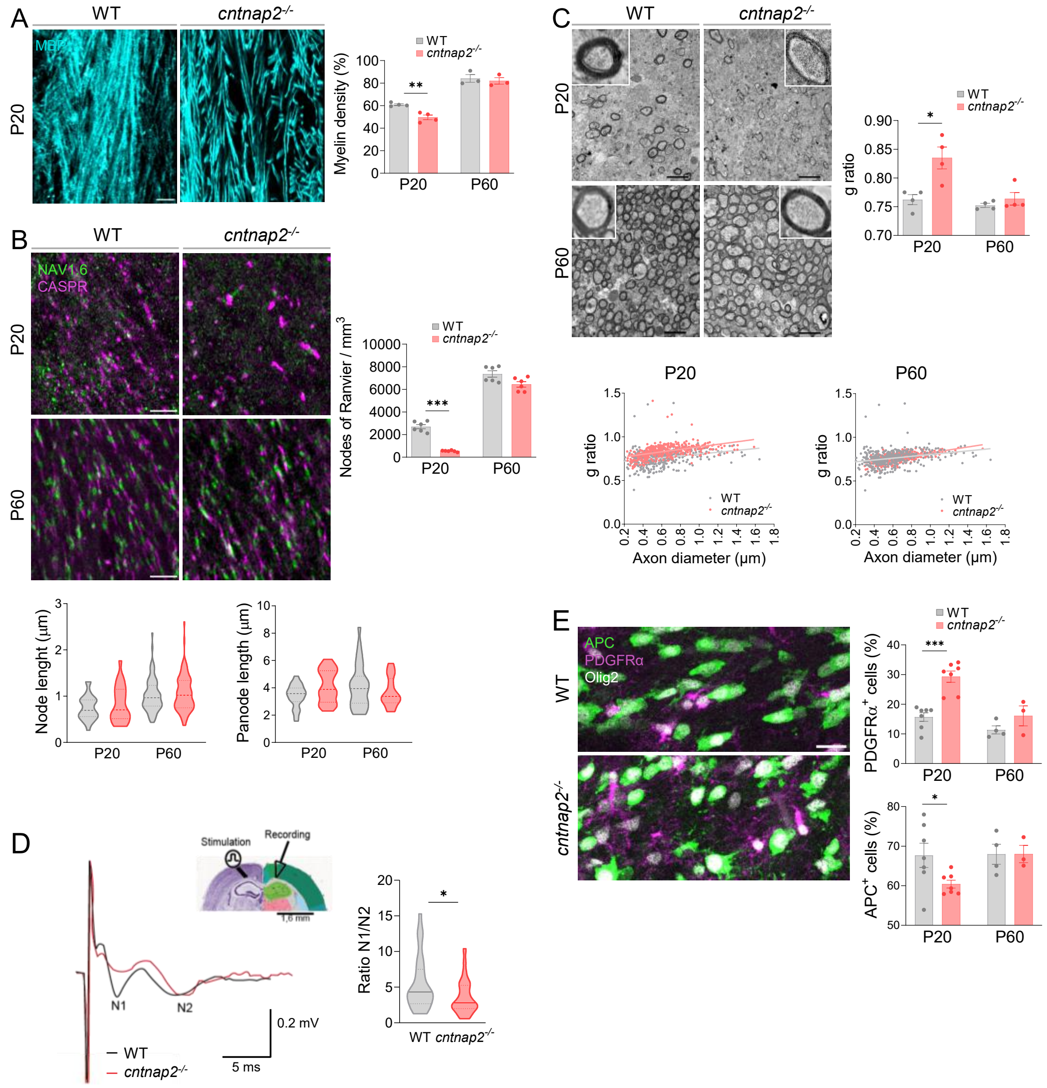


**Fig. S1.** *Cntnap2*$^{-/-}$ mice showed a decrease in myelination during development in corpus callosum. **A, B**) Confocal z stack images and quantifications of MBP coverage (A, n=4/4 at P20 and 3/3 at P60) and nodes of Ranvier identified by Nav1.6/ Caspr labeling (B, n=6 per group). **C**) TEM images of myelinated axons in the corpus callosum of juvenile WT and *cntnap2*$^{-/-}$ mice. Scale bar = 2 µm. G-ratio distribution (bottom) and g-ratio average (right) in WT and *cntnap2*$^{-/-}$ mice (n=4 mice per group). **D**) CAP recordings corresponding to N1 myelinated and N2 unmyelinated fibers in the corpus callosum of juvenile WT and *cntnap2*$^{-/-}$ mice. Histogram represents the ratio of N1 and N2 amplitudes (average of 12-15 recordings coming from n=3-5 mice). **E**) Confocal z stack images displaying triple staining, APC (oligodendrocytes), PDGFRα (OPCs) and olig2 in corpus callosum of juvenile WT and *cntnap2*$^{-/-}$ mice. Scale bar = 20 µm. Right, proportion of PDGFRα$^+$ OPCs and of APC$^+$ mature oligodendrocytes normalized to the total number of oligo2$^+$ oligodendrocytes in juvenile and adult WT and *cntnap2*$^{-/-}$ mice (n= 4/4 at P20 and 3/3 at P60). Data is represented as mean ± SEM. Statistical analysis were performed with two-tailed unpaired Student t-test. *p<0.05, ***p<0.001.

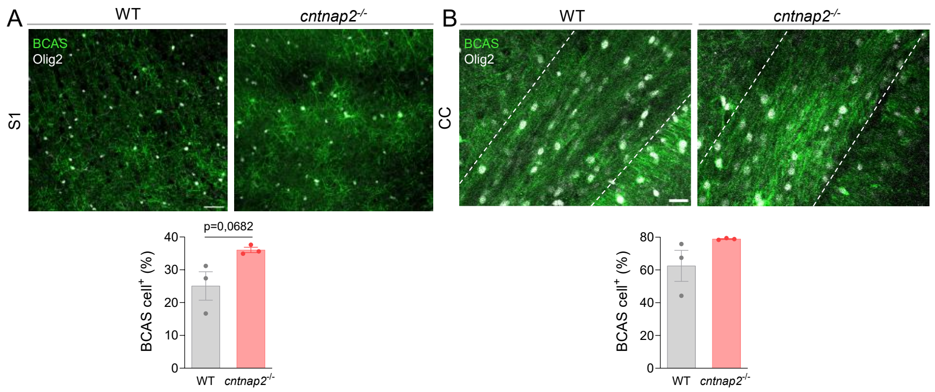


**Fig. S2.** Confocal z stack images and quantifications of BACS⁺ immature oligodendrocytes in layer V S1 and corpus callosum in juvenile WT and *cntnap2*$^{-/-}$ mice (n= 3/3). Scale bar = 50 μm (S1) and 25 μm (cc). Data is represented as mean ± SEM. Statistical analysis were performed with two-tailed unpaired Student t-test.

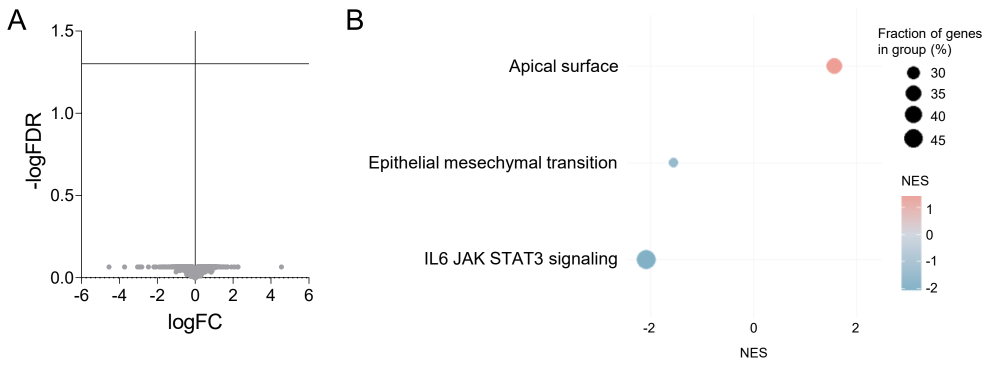


**Fig. S3**. Transcriptomic analysis in adult WT and *cntnap2*$^{-/-}$ oligodendrocytes. **A**) Volcano plot showing the absence of differentially expressed genes (DEGs; FDR q-value < 0.05) in adult *cntnap2*$^{-/-}$ mice. **B**) Hallmark pathway analysis of RNA sequencing data with Gene Set Enrichment Analysis (GSEA) showing the upregulated (red) and downregulated (blue) biological processes, ranked by normalized enrichment score (NES).

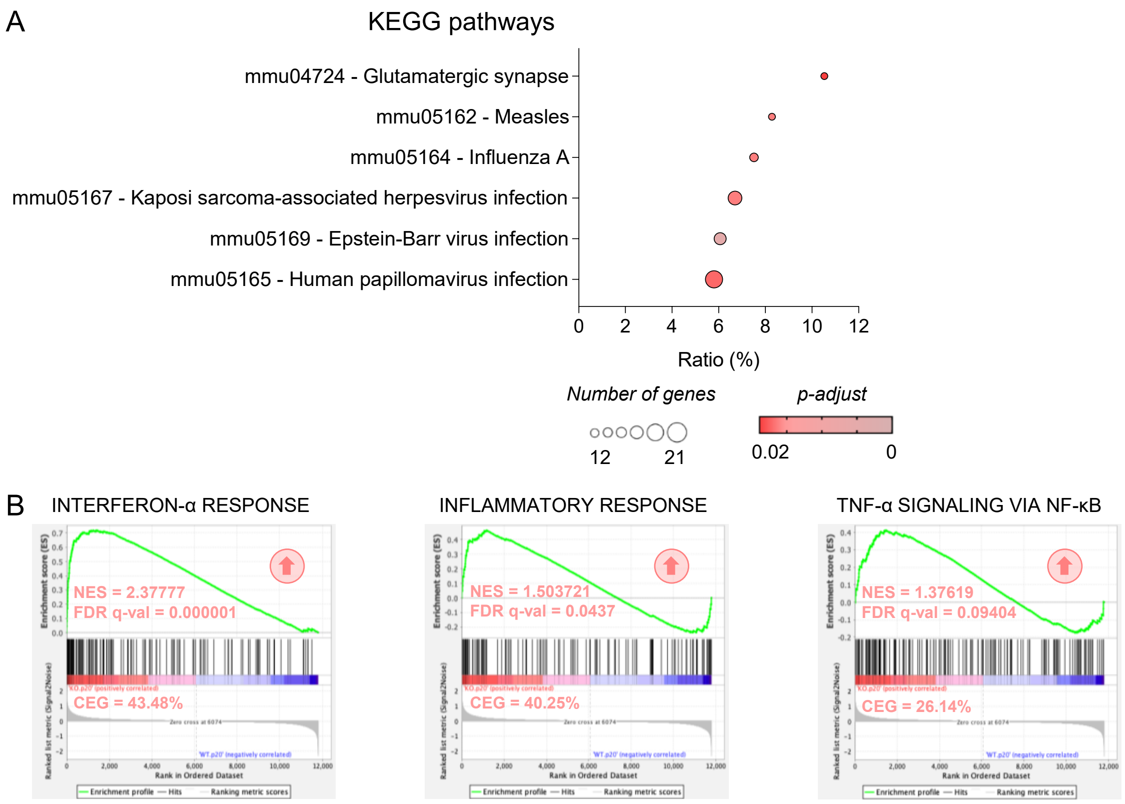


**Fig. S4**. Kyoto Encyclopedia of Genes and Genomes (KEGG) and GSEA analysis in oligodendrocytes derived from juvenile WT and *cntnap2*$^{-/-}$ mice. A) Top 6 biological pathways obtained by KEGG enrichment analysis, ranked from higher to lower ratio. This ratio indicates the percentage of significant genes over the total genes in a given pathway, colour spectrum ranges significance and dot size is proportional to the number of genes in each cluster. B) Enrichment plots (left) of Interferon a, inflammatory response and tumor necrosis factor gene sets from Hallmark analysis with corresponding NES, FDR q-value and Core Enrichment Genes (CEG).

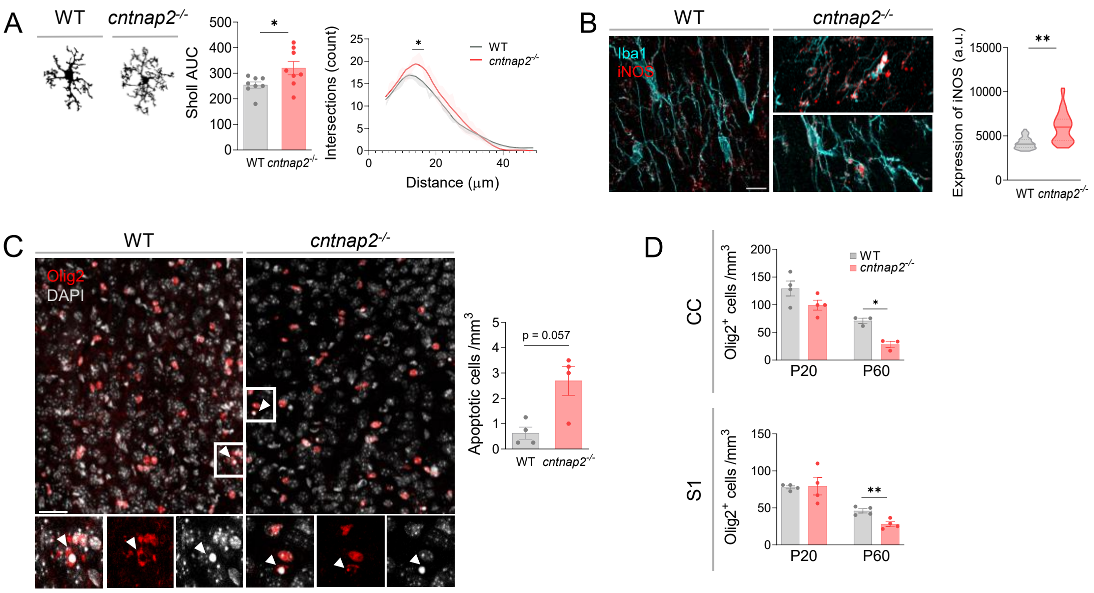


**Fig. S5.** Microgliosis in juvenile *cntnap2-/-* mice. **A**) Sholl analysis representing microglia complexity in S1 of juvenile WT and *cntnap2-/-* mice. **B**) Representative images and quantification of iNOS expression in the corpus callosum of juvenile WT and *cntnap2-/-* mice (n= 44 cells from 3 different mice). Scale bar= 10 μm. **C)** Confocal z stack images and quantification of apoptotic oligodendrocytes showing nuclei condensation in S1 of juvenile *cntnap2-/-* mice (n= 4). Scale bar = 50 μm. **D**) Quantification of the total number of Olig2+ oligodendrocytes at P20 and P60 in both corpus callosum and S1 (n =4). Data is represented as mean ± SEM. Statistical analysis were performed with 2-tailed paired student (A) and unpaired Student t-test (B-D). *p<0.05, **p<0.01.

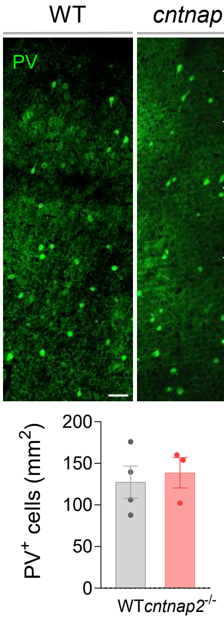


**Sup. Fig. 6.** Confocal images and analysis of the total number of PV+ interneurons in the S1 of adult WT and *cntnap2-/-* mice (n=4 per group). Scale bar = 50 μm.

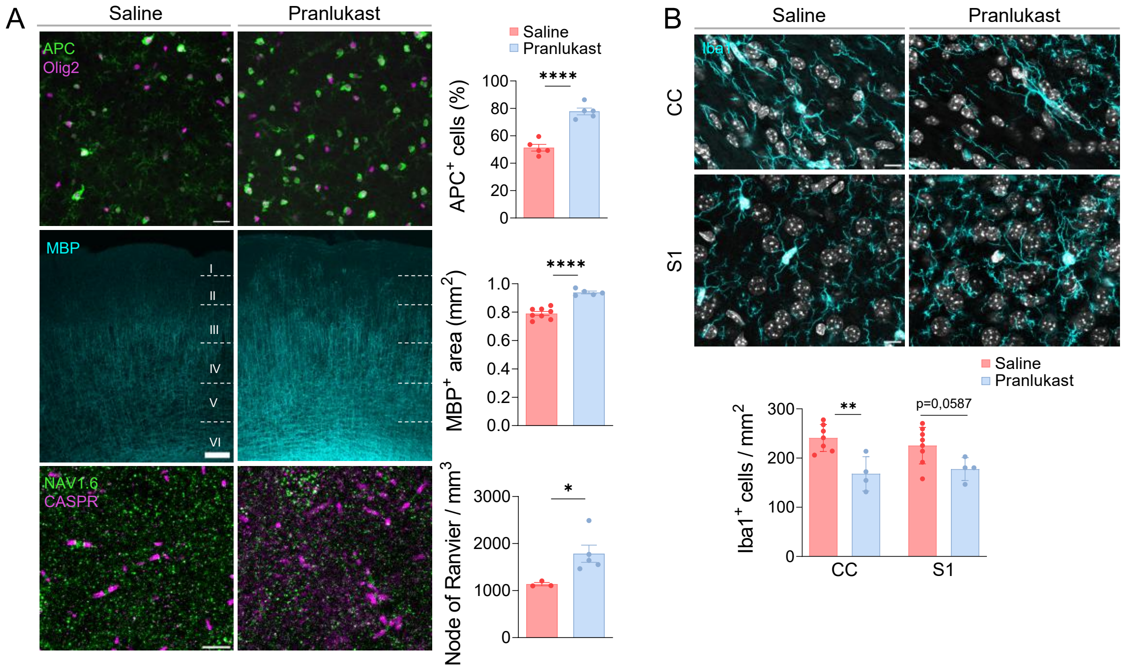


**Sup. Fig. 7**. Treatment with pranlukast, antagonist of the orphan receptor GPR17, reverses myelin deficits during development in *cntnap2*$^{-/-}$ mice. **A**) Confocal z stack images and quantifications of the proportion of mature APC$^+$ OLs (green) normalized to total Olig2 OLs (magenta) (top, n= 5), the S1 area myelinated (middle, n= 8/6) and the number of nodes of Ranvier (bottom, n= 3/5) in *cntnap2*$^{-/-}$ mice treated with saline or pranlukast. Scale bar = 25 μm (top), 50 μm (middle) and 5 μm (bottom). **B**) Confocal z stack images and quantification of microglial cells numbers in the corpus callosum and layer V of S1 in *cntnap2*$^{-/-}$ mice treated with saline or pranlukast (n = 7/4). Scale bar = 10 μm. Data is represented as mean ± SEM. Statistical analysis were performed with 2-tailed unpaired Student t-test (B-D). *p<0.05, **p<0.01, ****p<0,0001.

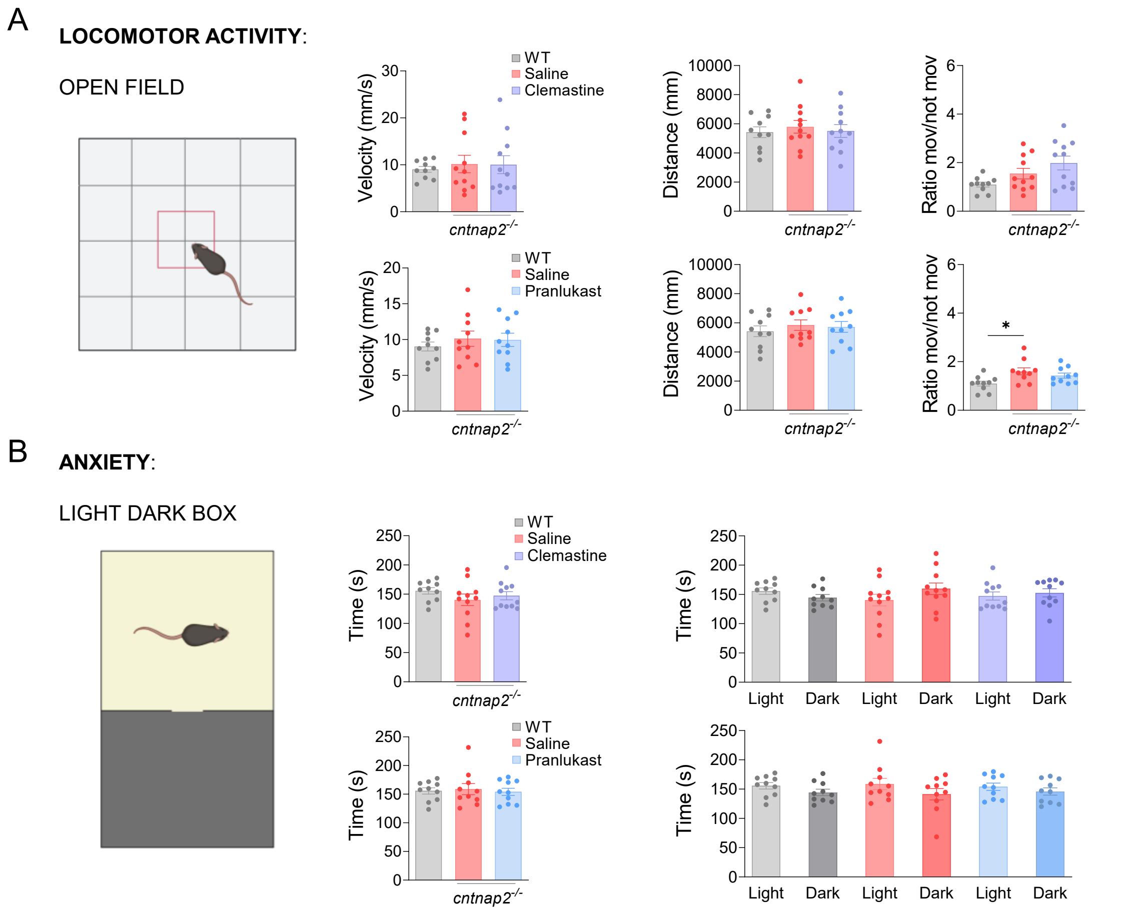


**Sup. Fig. 8.** Juvenile *cntnap2*$^{-/-}$ did not show any sign of anxiety. **A**) Velocity, distance travelled and time spent in movement or immobilized in WT mice and *cntnap2*$^{-/-}$ mice treated with saline, clemastine or pranlukast (n = 10-11). **B**) Time spent in light/dark (left) and time spent in each compartment WT mice and *cntnap2*$^{-/-}$ mice treated with saline, clemastine or pranlukast (n= 10-11). Data is represented as mean ± SEM. Statistical analysis were performed with ANOVA and Tukey's multiple comparison post-hoc test.
